# Enabling the prediction of phage receptor specificity from genome data

**DOI:** 10.64898/2026.04.02.716166

**Authors:** Lucas Moriniere, Avery J. C. Noonan, Alexey Kazakov, Melina Pena, Madeline Svab, Edwin O. Rivera-Lopez, Flavien Maucourt, Milo S. Johnson, Simon Roux, Britt Koskella, Adam M. Deutschbauer, Edward G. Dudley, Vivek K. Mutalik, Adam P. Arkin

**Affiliations:** California Institute for Quantitative Biosciences, University of California Berkeley, Berkeley, CA, USA; Environmental Genomics and Systems Biology Division, Lawrence Berkeley National Laboratory, Berkeley, CA, USA; Department of Molecular and Cell Biology, University of California Berkeley, Berkeley, CA, USA; Plant and Microbial Biology Department, University of California Berkeley, Berkeley, CA, USA; Department of Food Science, The Pennsylvania State University, University Park, PA, USA; Joint Genome Institute, Department of Energy, Berkeley, CA, USA; Department of Integrative Biology, University of California Berkeley, Berkeley, CA, USA; Chan-Zuckerberg San Francisco Biohub, San Francisco, CA, USA; Escherichia coli Reference Center, The Pennsylvania State University, University Park, PA, USA; Department of Bioengineering, University of California Berkeley, Berkeley, CA, USA

## Abstract

Predicting which receptor a phage binds to from genome sequence alone has remained an intractable challenge, principally because the experimental phenotypic data required to train and validate predictive models have not been available at sufficient scale. Here we address this by conducting 1,050 genome-wide genetic screens across 255 taxonomically diverse *Escherichia coli* dsDNA phages, assigning host receptors to 193 phages across 19 receptor classes. Comparative genomics and AlphaFold3 structural modelling resolved the sequence determinants of specificity to defined receptor-binding protein domains and individual residues. Machine learning models trained on this dataset predicted host receptor identity from phage genome sequence alone without prior annotation of receptor-binding genes, achieving perfect precision and greater than 80% recall on 49 independently validated phages, and yielding predictions for 1,060 of 1,875 *E. coli* phage genomes in NCBI. Domain swaps redirected receptor specificity as predicted, and a single amino acid substitution proved both necessary and sufficient to switch recognition between two distinct porins. These results demonstrate that systematic phenotyping at scale makes sequence-based prediction of molecular interaction specificity tractable, with direct implications for phage-based medicine, microbiome engineering and the broader challenge of inferring host-pathogen interaction outcomes from sequence.

## Introduction

Specific and selective molecular recognition underlies the organization of living systems at every scale, from immune surveillance^1,2^ and cell-cell communication^3^ to the assembly of multiprotein complexes^4^ and the specificity of host-pathogen interactions^5^. Predicting these interactions from sequence alone is an actively pursued goal in biology^6,7^, yet for most classes of molecular interactions, and most acutely for host-pathogen recognition, the experimental foundation needed to make interaction specificity predictable from sequence does not exist^8^.

Phage-bacteria interactions pervade every microbial ecosystem on Earth, driving bacterial evolution, mediating horizontal gene transfer, and regulating the composition and function of microbial communities from the human gut to the open ocean^9^. At the molecular level, each interaction is initiated by the recognition of a bacterial surface receptor by a phage receptor-binding protein (RBP), and it is this specificity that likely determines host range^10–14^, governs coevolution^15,16^, and defines the therapeutic potential of phage candidates^17^. The sequence determinants of this recognition remain uncharacterized for the overwhelming majority of phages. Among *Escherichia coli* double-stranded DNA (dsDNA) phages, the most intensively studied group in the field, experimentally validated receptor information is available for fewer than 200 phages^18–20^, and the receptor-binding proteins responsible for specificity have been characterized for a handful of model phages only^21^. The systematic characterization of receptor specificity across large, diverse phage collections has primarily been limited by experimental throughput. Barcoded genome-wide screening technologies, principally RB-TnSeq (random bar code transposon-site sequencing) for loss-of-function^22^ and Dub-seq (dual-barcoded shotgun expression library sequencing) for overexpression^23^, can resolve the full host genetic landscape of phage infection in a single experiment and are parallelizable across hundreds of phages without proportional cost increases. Prior applications to phage-host specificity have been limited to approximately 30 phages infecting *Enterobacteriaceae*^24–27^, a sample insufficient in scale and taxonomic diversity to support generalizable predictive modelling.

Here we close this gap by establishing a genotype-to-phenotype framework (**Figure 1A**) linking phage genome sequence directly to host receptor identity, trained and validated across a taxonomically comprehensive collection of *E. coli* dsDNA phages. This collection comprises 255 phages spanning nearly all major genomic clusters in current phage sequence space. Through 1,050 genome-wide screens, experimental receptor assignments were obtained for 193 phages across 19 independent receptor classes. Comparative genomics traced the sequence determinants of specificity to defined receptor-binding protein domains, and AlphaFold3 structural modelling localized these determinants to binding interfaces. Machine learning models trained on this dataset predict receptor identity from phage genome sequence alone without prior annotation of receptor-binding genes, yielding predictions for 1,060 of the 1,875 *E. coli* phage genomes currently available in NCBI. Receptor-switching experiments informed by these predictions, including bidirectional single-residue mutagenesis, confirm that the identified sequence features are causal determinants of receptor recognition and demonstrate that receptor identity can be rationally reprogrammed.

**Figure 1.**
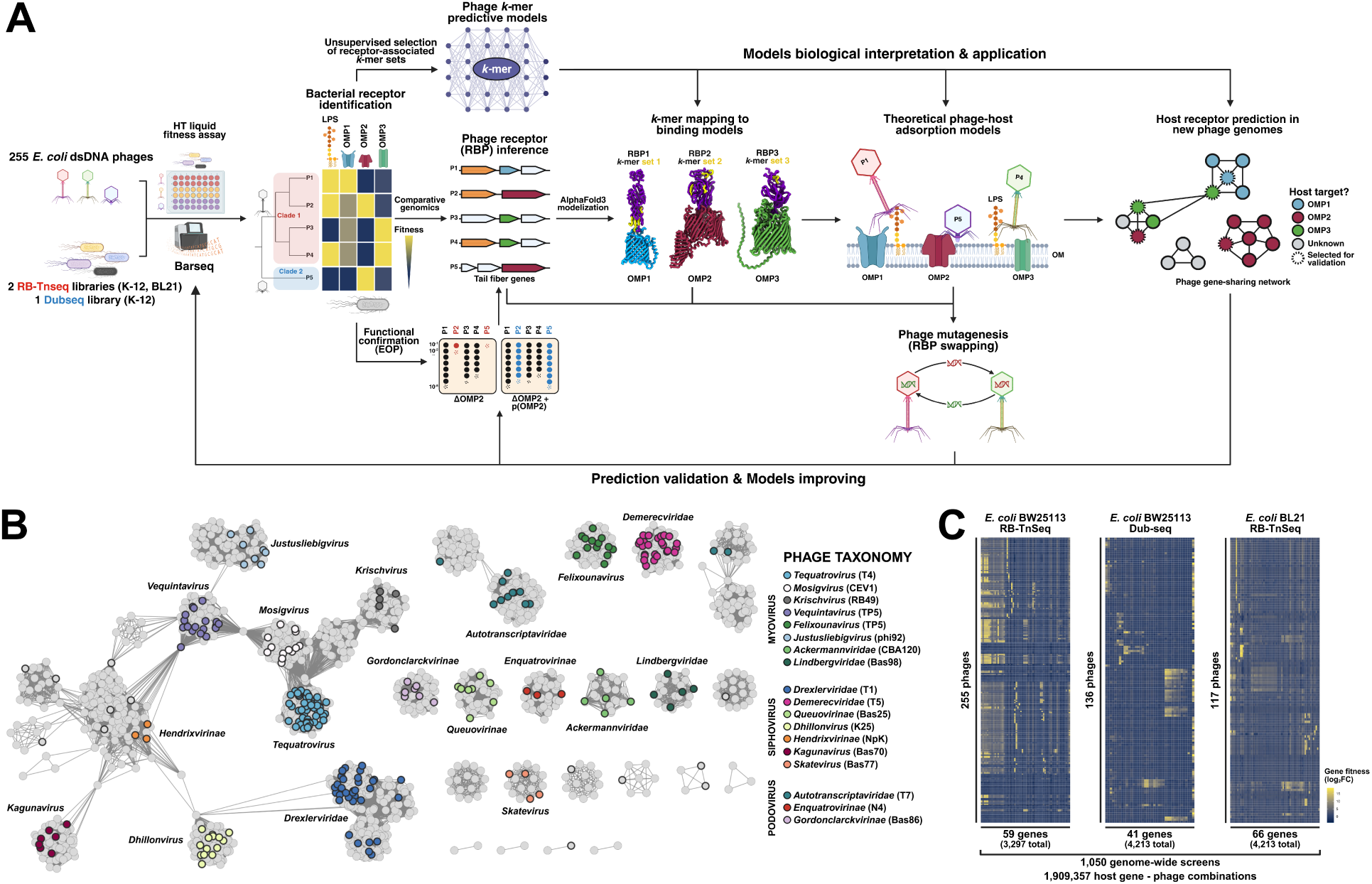
Overview of the experimental workflow, biological resources, and genome-wide-screen datasets generated. (**A**) Schematic of the experimental and computational workflow developed to characterize and predict phage receptor specificity from genome data. A collection of 255 phages has been tested in high-throughput liquid assays on 3 genome-wide libraries. Results were interpreted to assign one or multiple host receptors to each phage, and subsets of readouts were validated by determining the EOP on single-gene deletion or overexpression mutants. Phenotype-informed comparative genomics was then used to identify putative receptor-binding protein (RBP) genes in the phage genomes, while an unsupervised machine learning-based strategy was applied to identify predictive *k*-mer features in phage proteomes associated with specificity towards 13 of the 19 receptor classes. Interactions between the putative RBPs and outer membrane protein (OMP) host receptors were modelled with AlphaFold3 and predictive *k*-mers were mapped on cognate RBP-receptor pairs. Receptors were predicted on phage genomes retrieved from the NCBI database, and a subset of phages were acquired to experimentally validate the predictions and improve the models. Receptor-switching experiments were conducted by swapping RBPs between phages to demonstrate causality and show that receptor specificity can be rationally reprogrammed. (**B**) vConTACT2 gene-sharing network of 1,875 *E. coli* dsDNA phages from the NCBI GenBank database and the 255 phages in our collection. Phages used in this study are circled in black, and colored by their taxonomy when relevant. Examples of representative phages for each taxon are indicated in the legend. (**C**) Complete heatmaps of every high-scoring gene uncovered in the RB-TnSeq and Dub-seq phage experiments. Numbers of phages tested, high-scoring genes uncovered, and total number of genes assayed in each genome-wide library are indicated.

## Results

### A diverse collection of 255 E. coli dsDNA phages spanning near-complete sequence space

To generate a phenotypic dataset of receptor specificity sufficient for predictive modelling, we assembled and verified a collection of 255 unique *E. coli* dsDNA phages (**Table S1**), overwhelmingly lytic and all confirmed by whole-genome (re-)sequencing. We primarily sought contributions from the broad research community, allowing us to gather 224 phages isolated over a century and for diverse purposes (see **Methods - Phage acquisition and amplification**). An additional 31 phages were isolated in-house using diverse *E. coli* strains (**Table S2**). The three morphotypes within the *Caudoviricetes* class are represented: 129 myoviruses, 101 siphoviruses, and 25 podoviruses, spanning at least 10 families, 19 subfamilies, and 50 phage genera^28^. To assess the diversity of this collection relative to known *E. coli* phage sequence space, we performed a gene-sharing network analysis^29^ of 1,875 available tailed (*Caudoviricetes* class) *E. coli* phage genomes in the NCBI GenBank database. Our 255 phages span nearly all genomic clusters identified in this analysis (**Figure 1B**), with proteomic equivalence quotients (PEQ)^30^ confirming that isolates within clades were non-redundant and non-clonal (**Figure S1**). Commonly isolated clades were overrepresented^18^, including 73 T-even-like (*Straboviridae*) and 27 rv5-like (*Vequintavirinae* and *Justusliebigvirus*) myoviruses, 64 T1-like (*Drexlerviridae* and related taxa *Queuovirinae*, *Dhillonvirus* and *Hendrixvirinae*) and 25 T5-like (*Demerecviridae*) siphoviruses, and 11 T7-like (*Studiervirinae*) podoviruses, consistent with their ecological prevalence and their practical importance as candidate therapeutic and biotechnological agents.

To facilitate the exploration of our phage collection, we created the Phage Datasheets (https://iseq.lbl.gov/PhageDataSheets/Ecoli_phages/), an interactive web-based platform that centralizes the data generated for each phage in this study. In addition, phage genomes can be visualized and investigated using a variety of comparative genomics tools through the interconnected Phage Genome Browser (https://iseq.lbl.gov/ecoliphages).

### 1,050 genome-wide screens assign host receptors to 193 phages across 19 receptor classes

Leveraging the scalability of barcoded genome-wide screens, we challenged RB-TnSeq and Dub-seq libraries in *E. coli* BW25113 and BL21 with the full phage collection (**Figure 1C**). These two strains lack O-antigen (OPS) and capsular (CPS) polysaccharides, allowing direct probing of outer membrane protein (OMP) receptors for phages that do not require prior recognition of these surface structures. All 255 phages were screened on the BW25113 RB-TnSeq library (3,728 genes covered), yielding 59 high-scoring disrupted genes (**Dataset S1**). A subset of 136 phages was additionally tested with the BW25113 Dub-seq library (4,213 genes covered), identifying 41 high-scoring overexpressed genes (**Dataset S2**). Screens on the BL21 RB-TnSeq library (3,297 genes covered) were conducted with 117 phages, yielding 66 high-scoring genes (**Dataset S3**), yet they essentially mirrored BW25113 results and were deprioritized. In total, and accounting for biological replicates, 1,050 genome-wide screens were conducted (763 RB-Tnseq and 287 Dub-seq individual experiments), collectively testing over 1.9 million unique gene-phage combinations. Subsets of high-scoring genes were cross-validated by efficiency-of-plating (EOP) assays on BW25113 single-gene deletion or overexpression mutants^31,32^, comprising more than 2,000 individual observations (**Datasets S4-S5**). These targeted assays were particularly valuable for validating LPS biosynthesis genes and for identifying phages targeting the essential transporter LptD, the latter of which cannot be assessed by transposon insertion. The complete fitness data generated in the genome-wide screens can be visualized and explored through a BarSeq browser implemented in the Phage Datasheets.

A specific host receptor or combination of receptors was assigned to 193 of 255 phages based on improved survival rates for cells presenting mutations in the corresponding genes (**Table S1, Datasets S1-S3**). Of the 62 remaining phages, 31 were previously demonstrated to require the O16-antigen for infection^19^, another 23 belong to clades known to recognize OPS or LPS moieties, including *Felixounavirus, Ackermannviridae* and *Autotranscriptaviridae*^18,33^, 4 are temperate phages exerting insufficient selective pressure on the mutant pools to yield interpretable screen readouts (NpA, NpB, NpD & NpO), 3 are hypothesized to require the expression of a specific capsule for infection (HSDP1, HSDP2 & HSDP3), and one putatively recognizes an unknown OMP absent from BW25113 (Mnm3).

We phenotypically resolved receptor identity into 19 distinct receptor classes, including specific outer membrane proteins (Tsx, FadL, OmpA, OmpF, OmpW, OmpC, FhuA, BtuB, LptD, NupG, LamB, TolC, YncD), LPS core sugar moieties (Kdo, HepI, HepII, GluI, HepIV) and the N4-glycan receptor polysaccharide (NGR). For downstream modelling, 13 receptor classes are defined operationally as receptor targets represented by at least 4 phages in the training set, each corresponding to one binary classifier: 8 OMP receptors (Tsx, OmpA, OmpF, OmpC, FhuA, BtuB, LptD, LamB), 4 LPS core sugars (Kdo, HepI, HepII, GluI) and NGR (**Table S3**). Within the 73 *Straboviridae* phages, screens resolved both primary (porin) and secondary (LPS core sugar) receptors, assigning receptors to 71 of 73 phages (**Figure 2A**). Five known porins were confirmed as primary receptors (Tsx, FadL, OmpA, OmpF, OmpC), and OmpW was identified as the primary receptor of phage RB68, the first known *E. coli* phage to target this small porin^34^. Among secondary receptors, 40 phages displayed a T4-like phenotype relying on Kdo, 29 required HepI, and cross-validation experiments confirmed that phages Ox4, TP7 and TulB additionally depend on GluI (**Figure S2**).

**Figure 2.**
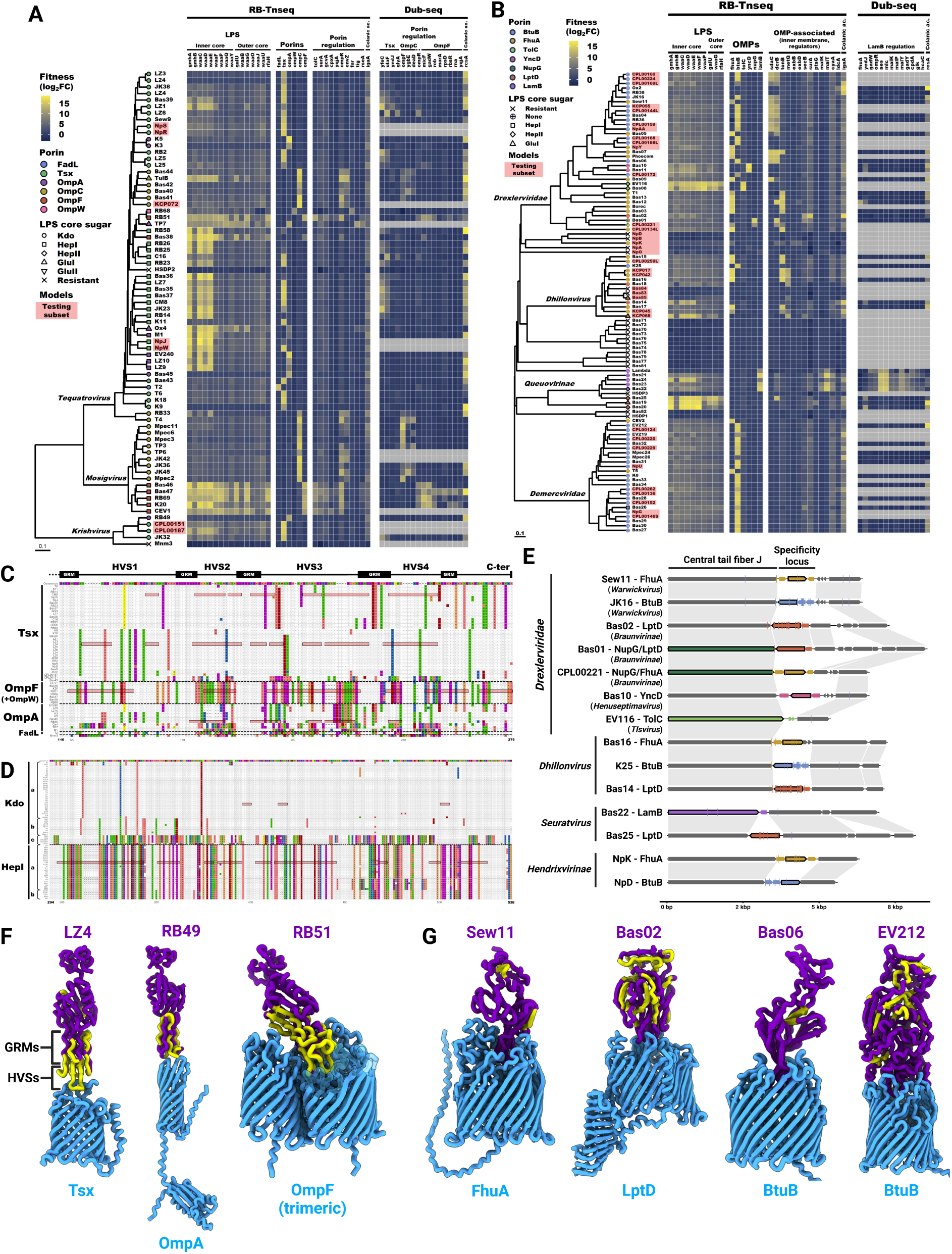
Host receptor and phage RBP identification in (**A, C-D, F**) *Straboviridae* and (**B, E, G**) siphovirus phages. (**A-B**) Annotated heatmaps showing the high-scoring genes, related membrane structures, and the resulting receptor(s) assignment in (**A**) 73 *Straboviridae* phages and (**B**) 101 siphovirus phages. Phages included in the model testing subset are highlighted in red. Grey boxes indicate missing data. (**C-D**) *Straboviridae* phage RBP C-terminal sequence alignments displaying amino acid variations from the consensus for (**C**) 54 Gp38 adhesins targeting Tsx, OmpF, OmpW, OmpA or FadL and (**D**) 73 Gp12 short tail fibers targeting the Kdo or HepI LPS core sugars. Contiguous predictive features are highlighted in red for representative phages. In (**C**), GRM = glycine-rich motifs, and HVS = hypervariable sequences. In (**D**), phage genera are indicated by the following: a = *Tequatrovirus*; b = *Mosigvirus*; c = *Krischvirus*. (**E**) Synteny plot of the central tail fiber J and downstream variable receptor specificity locus for representative *Drexlerviridae* and related phages. Receptor-specific genes are colored according to their target, and *bona fide* RBPs are highlighted by a black border. Location of singular predictive features is indicated on the gene tracks by colored dashes. Grey homology boxes linking gene tracks correspond to > 30% amino acid identity over > 50 % coverage. (**F-G**) Representative OMP-RBP interaction models generated with AlphaFold3 for (**F**) 3 *Straboviridae* phages targeting Tsx (LZ4), OmpA (RB49) and OmpF (RB51), and (**G**) 4 siphovirus phages targeting FhuA (Sew11), LptD (Bas02) and BtuB (Bas06 and EV212). Bottom blue proteins are the host OMPs, and top purple proteins are the phage RBPs. Structural location of contiguous predictive features is indicated in yellow on the phage RBPs.

Among the 101 siphoviruses, receptors were attributed to 81 phages (**Figure 2B**). Screens identified six known OMPs (FhuA, BtuB, LptD, LamB, TolC, YncD) and a novel receptor, the nucleoside transporter NupG, targeted by *Braunvirinae* phages Bas01 and CPL00221. These two phages exhibited dual-receptor phenotypes, with Bas01 requiring both NupG and LptD, whereas CPL00221 could infect Δ*nupG* and Δ*fhuA* mutants with approximately 10,000-fold reduced efficiency (**Figure S3**). Taxon-specific requirements for inner membrane or periplasmic proteins (DcrB, SdaC, MetQ) in defined siphovirus subgroups likely reflect post-adsorption steps rather than initial receptor recognition. Conclusive screens of other myoviruses and podoviruses revealed targeting of the NGR polysaccharide or diverse LPS core sugars (**Figures S4-S5**).

### Comparative genomics identifies receptor-binding proteins and domain-level determinants of specificity

Having assigned host receptors to 193 phages, we used comparative genomics to identify the corresponding receptor-binding proteins. Phages targeting the same receptor were expected to share conserved RBP-encoding genes or domains, with synteny breaks in tail gene regions marking receptor specificity loci. Candidate RBP sequences and loci were systematically aligned to identify specificity-associated domains^35^ (**Figure 2C-E**), and AlphaFold3 interaction modelling was used to identify the most structurally compatible RBP-OMP pairing where multiple candidates existed (**Figures 2F-G**).

In the *Straboviridae* family, receptor-associated sequence variation in Gp38 adhesins had previously been examined in smaller phage subsets^20,36,37^. Here, sequences of 54 Gp38 adhesins specifically recognizing one of five host porins (Tsx, FadL, OmpA, OmpF and OmpW) were retrieved and aligned (**Figure 2C**). Four C-terminal hypervariable sequence domains (HVSs) interspersed with conserved glycine-rich motifs (GRMs) were identified, consistent with the modular architecture described by Trojet et al.^36^. Allelic variability in the HVSs was specifically and consistently associated with porin targeting across all subgroups, with distinct alleles within each porin-specific subgroup potentially reflecting binding to different outer loop variants of the same porin^20^. The unique OmpW specificity phenotype displayed by phage RB68 seemed to be associated with a single amino acid change in its Gp38 variant’s HVS3 loop (**Table S4**), suggesting that point mutation in Gp38’s binding interfaces could be enough to alter receptor specificity. In the five phages of the *Krishvirus* genus, *gp38* was located approximately 55 kb downstream of *gp37* rather than in its canonical adjacent position, suggesting displacement or independent acquisition by horizontal gene transfer in the last common ancestor of this genus. Alignment of 73 Gp12 short tail fiber sequences confirmed that C-terminal allelic variability correlated with LPS core sugar targeting^18^, Kdo or HepI, and with phage genera (**Figure 2D**), indicating that both primary and secondary adsorption determinants are organized in discrete exchangeable sequence modules.

In the *Drexlerviridae* family and related taxa, we confirmed prior observations^18^ that a hypervariable locus directly downstream of the central tail fiber gene *gpJ* systematically correlates with receptor specificity (**Figure 2E**). Four unrelated protein families encoded within this locus were each specifically associated with targeting of FhuA, BtuB, LptD or YncD, defined by more than 30% amino acid identity and more than 99% coverage within each family, and AlphaFold3 interaction models supported their roles as *bona fide* RBPs (**Figure 2G**). These putative RBPs were frequently co-localized with one or two accessory proteins likely involved in host specificity determination. CryoEM experiments recently demonstrated that the *bona fide* RBP of phage Bas18 was indeed binding to the extracellular face of LptD, and that the adjacent accessory protein of phage Rtp was a superinfection exclusion factor binding to the periplasmic face of LptD^38^. In parallel, distal domains of the central tail fiber GpJ itself were associated with targeting of LamB, TolC^18^ or NupG in distinct phage subsets (**Figures S6-S7**), explaining the dual OMP requirements of Bas01 and CPL00221 through co-occurrence of a NupG-specific GpJ domain alongside a locus-encoded FhuA- or LptD-specific RBP. This comparative genomics approach additionally enabled computational receptor attribution for eight phages lacking clear screen phenotypes, with locus structure confidently indicating targeting of LptD (Bas83, Bas84 and HSDP3), YncD (NpA and NpB), BtuB (NpD), LamB (NpO), or OmpC (HSDP2). The broader pattern of conservation and synteny breaks at this locus is consistent with horizontal gene transfer as the primary driver of receptor specificity diversification in *Drexlerviridae* and related taxa.

In the *Demerecviridae* family, FhuA and BtuB recognition by T5-like siphoviruses represents an independent evolutionary origin relative to *Drexlerviridae*, with structurally distinct RBPs occupying the same genomic locus upstream of the terminase small subunit gene ( **Figure 2G**). FhuA recognition was restricted to T5 and CEV2, both encoding the same RBP (*pb5* in T5^39^), while the 23 BtuB-binding *Demerecviridae* phages carried a homologous but distinct protein sharing approximately 30% amino acid identity with pb5, suggesting divergence from a common ancestral RBP with subsequent receptor specificity change.

The RBPs targeting the NGR polysaccharide in myoviruses of the *Vequintavirus* and *Justusliebigvirus* genera were identified by sequence comparison with the known Gp65 tail sheath RBP of the N4 podovirus^40,41^. A small 59 to 76 amino acid-long domain of N4’s Gp65 showed significant homology, exceeding 30% amino acid identity, within the poly-GlcNac deacetylase domain^18^ of a tail fiber protein present in all NGR-binding myoviruses (**Figure S8**), exemplified by Gp142 in phage phi92^42^.

Across these families, receptor identity is encoded in modular sequence elements operating at multiple organizational scales, encompassing the exchange of entire RBP genes between loci (gene-scale modularity), allelic variation within conserved scaffolds (domain-scale modularity), and point mutation at predicted binding interfaces (residue-scale modularity). Taken together, RBPs were inferred for 200 phenotypically characterized phages, and 108 RBP-host OMP interactions were modelled with AlphaFold3 with sufficient confidence to support structural interpretation (**Tables S1, S5-S7**).

### Annotation-free models predict receptor identity from phage proteome sequence

To test whether receptor specificity could be predicted directly from phage proteome sequence without prior annotation of receptor-binding genes, we constructed genome-scale *k*-mer feature representations from complete phage proteomes using the GenoPHI framework^43^. For each of 13 receptor classes represented by at least 4 phages, independent binary classifiers were trained using recursive feature elimination and gradient-boosted decision trees (**Figure 3A**). Model performance was evaluated by 20-fold nested cross-validation with balanced 10% holdouts, restricting feature selection to training partitions at each fold. Models achieved a median AUROC of 0.99 (range 0.62–1.00) and a median AUPR of 0.95 (range 0.23–1.00) (**Figure 3B, Figure S9, Table S3**). Models were trained and predictive features extracted independently of the comparative genomics analysis described above, the convergence reported below was established *post hoc*.

**Figure 3.**
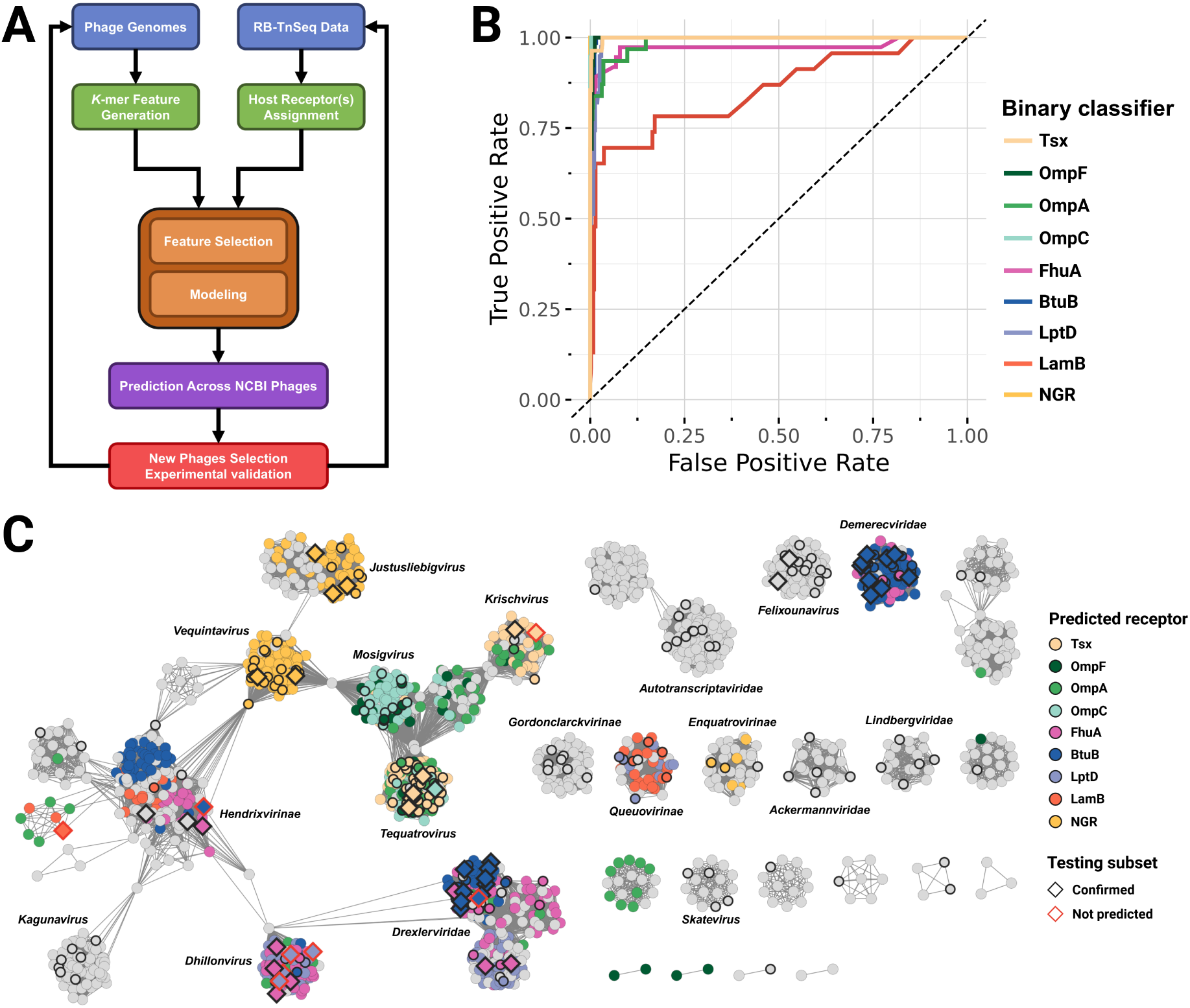
Modeling strategy and predictions across 1,875 NCBI *E. coli* phages. (**A**) Schematic of the iterative modeling strategy. An initial set of 206 phage proteomes was split into *k*-mer features and host receptor(s) were assigned to phages based on RB-TnSeq data. Unsupervised modeling of 13 independent binary classifiers selected predictive features for each receptor class. Predictions were made in 18,398 *Caudoviricetes* phage genomes from the NCBI GenBank database, and a testing subset of 49 new phages was acquired to experimentally validate the predictions. The resulting RB-TnSeq and genome data were used to re-train the final models on a set of 255 phages. (**B**) Receiver operating characteristic (ROC) curves showing the sensitivity (true positive rate) and specificity (true negative rate) of the 8 OMP-specific and the NGR-specific binary classifiers. (**C**) vConTACT2 gene-sharing network displaying predictions of 9 binary classifiers (8 OMPs and NGR) across 1,875 *E. coli* dsDNA phages from the NCBI GenBank database and the 255 phages in our collection. Phages included in the initial training set (206 phages) are represented with a circle, while phages selected for the testing subset (49 phages) are indicated by a diamond. Colors indicate the predicted receptor for NCBI phages or experimentally determined receptor for phages in the training and testing subsets.

The *k*-mer features identified by these models (**Dataset S6**) were mapped against the receptor-binding domains resolved by comparative genomics, providing an independent test of whether the predictive signatures correspond to genuine molecular determinants of recognition. In *Straboviridae*, porin-specific *k*-mers were detected in Gp37 and Gp38 (**Figure 2C, Figure S10)**, and LPS sugar-specific features in Gp12 (**Figure 2D, Figure S11**), the upstream fibritin neck whisker and downstream baseplate wedge subunit genes. In Gp38 adhesins, predictive *k*-mers localized to the C-terminal GRMs and HVSs, and AlphaFold3 interaction models confirmed their overlap with exposed binding loops that directly contact porin receptors^36^ (**Figure 2F**). For OmpC-targeting phages, *k*-mers were identified in the D10 and D11 C-terminal domains of Gp37^44,45^, with extensive allelic variation in the D11 “tip” subdomain (**Figure S10**) explaining why features localized to surrounding conserved regions rather than the known OmpC-binding residues of T4’s LTF. For siphovirus models covering FhuA, BtuB, LptD and LamB, *k*-mer features were detected in the putative RBPs and their co-localized accessory proteins (**Figure 2E**), though mapping onto AlphaFold3 interaction models showed that they highlighted core structural domains rather than direct binding interfaces, in contrast to the *Straboviridae* adhesins (**Figure 2G**). The models additionally identified the C-terminal tip domain of GpJ in *Drexlerviridae* and related taxa as a specificity-associated feature (**Figure S12**), suggesting it could play an important role in mediating structural compatibility between GpJ and the locus-encoded RBP. For the NGR-specific model, a perfectly conserved five-amino acid motif ‘GMSHY’ was detected within the small amino acid domain shared between NGR-targeting myoviruses and podoviruses (**Figure S8**), whose evolutionary distance and morphological differences suggest convergent evolution rather than horizontal gene transfer as the origin of their shared receptor specificity.

For 11 of 13 modeled receptor classes (all except HepII and GluI), predictive *k*-mer features mapped to the same receptor-binding proteins identified by comparative genomics and, for 6 receptor classes where pre-existing data or AlphaFold3 models were available (Tsx, OmpA, OmpF, Kdo, HepI, NGR), to predicted interface-proximal residues. This convergence from three orthogonal analyses (systematic phenotyping, comparative genomics and sequence-based prediction) supports the conclusion that these features are genuine determinants of receptor recognition rather than proxies for phylogeny or genomic context. In several instances, the sequence-based models exceeded the resolution achievable by comparative genomics alone, detecting specificity-associated features at the scale of individual amino acids, a level of granularity that alignment-based and structural approaches did not independently resolve.

### Iterative experimental validation confirms model precision across 49 independently tested phages

To validate model performance on genuinely novel phages, models were first trained on 206 experimentally characterized phages (**Table S1**) and applied to 18,398 *Caudoviricetes* genomes retrieved from the NCBI GenBank database (accessed in May 2025). Among these 18,398 genome sequences, 1,875 corresponded to *E. coli* phages, of which 49 were selected based on their availability and predicted receptor specificity, acquired, and experimentally characterized using the same genome-wide screening workflow applied to the training set (**Figures 2A-B, Figure S4**). Final model iterations trained on the complete 255-phage dataset were then applied to all 18,398 genomes, yielding 3,123 phages with at least one receptor prediction (**Dataset S7**).

At least one receptor was predicted for 1,060 of the 1,875 *E. coli* phage accessions in the NCBI GenBank database (56.5%), comprising 874 predictions for OMP receptors or NGR (**Figure 3C**) and 599 for LPS core sugars (**Figure S13**). Predictions were concentrated in taxa well represented in the training set, and their distribution across phage families was consistent with experimentally observed receptor diversity, for instance with FhuA and BtuB predictions restricted to siphoviruses and NGR predictions confined to *Vequintavirus*, *Justusliebigvirus* and *Enquatrovirus*. The OmpA model showed the highest rate of likely erroneous predictions in genomes lacking a Gp38 adhesin homolog, probably reflecting incomplete coverage of the extensive allelic variability within OmpA-binding adhesins in the 10-phage training panel (**Figure 2C**).

The 49-phage testing subset spanned 13 taxa, including two absent from training, the *Hendrixvirinae* temperate subfamily and the *Drexlerviridae* subfamily *Rogunavirinae*. Accurate predictions were made for OMP or NGR specificity in 41 of 49 phages (83.7%) and for LPS core sugar specificity in 47 phages (96%) (**Table S8**), with correct predictions achieved even in the two novel taxa. Where models failed, insufficient coverage of RBP sequence diversity in the training set was the limiting factor rather than a modelling limitation, as illustrated by the LptD model missing four *Dhillonvirus* phages because only two *Dhillonvirus* phages were present in training, and the Tsx model missing *Krishvirus* CPL00151 whose Gp38 subtype was represented by a single training phage (**Figure 2C**). No incorrect positive calls were observed among predictions that models made, implying perfect precision. Per-class precision and recall with exact 95% Clopper-Pearson confidence intervals are reported in **Table S3**.

### Rational reprogramming of receptor identity

Having identified receptor-associated sequence features computationally and localized them to specific receptor-binding modules, we tested whether these features represent causal determinants of receptor identity through targeted engineering in *Straboviridae* phages. We pursued three objectives: (a) complete reprogramming of both adsorption steps through RBP gene exchange, (b) identification of the minimal sequence changes sufficient to redirect specificity, and (c) prospective testing of whether the predictive models correctly anticipate the phenotypes of engineered variants.

Three T6-like phages with high proteomic similarity but distinct receptor specificity profiles were selected as the experimental system, specifically CM8 (Tsx/HepI), EV240 (OmpA/HepI) and JK38 (Tsx/Kdo) (**Figure 4A-B**). Replacing CM8’s *gp38* with that of EV240 generated CM8×EV240Gp38, whose RB-TnSeq profile was virtually indistinguishable from EV240, confirming a complete switch to OmpA/HepI specificity. Replacing CM8’s *gp12* with the JK38 variant generated CM8×JK38Gp12 with a Tsx/Kdo phenotype matching JK38. The double mutant CM8×JK38Gp12×EV240Gp38, constructed by introducing EV240’s *gp38* into CM8×JK38Gp12, displayed an OmpA/Kdo phenotype absent from all three parental phages, demonstrating that novel specificity combinations can be generated *de novo* by combinatorial RBP exchange. Whole-genome sequencing revealed that only the distal half of *gp38* from HVS2 onwards had been exchanged, confirming that the C-terminal hypervariable domains are the functional determinants of porin recognition.

**Figure 4.**
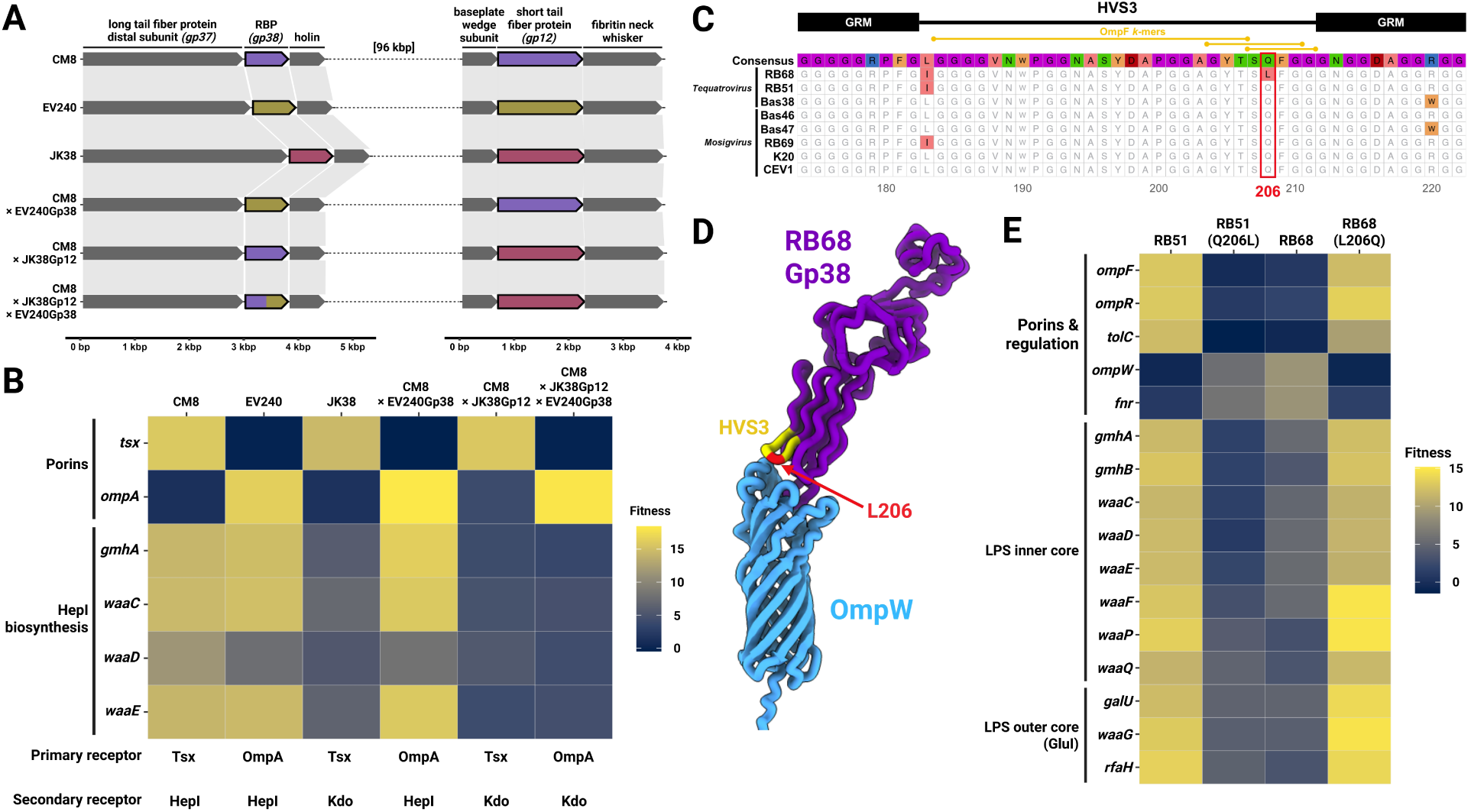
Receptor-switching experiments through RBP swapping in *Straboviridae* phages. (**A**) Schematic representation of the Gp38 and Gp12 variant combinations in the wild-type (CM8, EV240, JK38) and mutant phages. (**B**) High-scoring gene profiles obtained in RB-TnSeq experiments conducted on BW25113 with the above wild-type and mutant phages, showing expected changes in receptor specificity. (**C**) Localized Gp38 sequence alignment of domain HVS3 for all OmpF-specific phages and the unique OmpW-specific phage, RB68. Amino acid residues varying from the consensus are colored, and the causal receptor-switching Q206L mutation is highlighted by a red box. Overlapping OmpF-predictive *k*-mer features are shown above the alignment track. (**D**) AlphaFold3 interaction model of RB68’s Gp38 and OmpW highlighting the location of the Q206L mutation in the HVS3 domain (in yellow), at the interface with OmpW. (**E**) High-scoring gene profiles obtained in RB-TnSeq experiments conducted on BW25113 with the RB51 and RB68 wild-type and mutant phages, showing expected changes in receptor specificity.

Phage RB68, the first known *E. coli* phage targeting OmpW, provided a natural experiment in minimal sequence divergence. Its genome differed from that of its OmpF-targeting relative RB51 by only six polymorphisms (**Table S4**), among which a Q206L substitution in the HVS3 domain of Gp38 was uniquely absent from all other OmpF-binding phages in the collection (**Figure 4C**). This residue overlapped with *k*-mer features from the OmpF-specific model and was placed at the tip of a C-terminal binding loop by AlphaFold3 modelling (**Figure 4D**). Bidirectional mutagenesis confirmed causality: RB51(Q206L) switched from OmpF to OmpW recognition, while RB68(L206Q) reverted to OmpF specificity (**Figure 4E**), establishing that a single substitution at the binding interface is both necessary and sufficient to redirect specificity between two distinct porins.

The predictive models correctly anticipated the phenotype of every engineered variant (**Table S9**). CM8×EV240Gp38 received a correct OmpA prediction, CM8×JK38Gp12 and CM8×JK38Gp12×EV240Gp3 predictions for both HepI and Kdo, and the novel OmpA/Kdo combination in CM8×JK38Gp12×EV240Gp3 was correctly inferred from the swapped domains. For the single-substitution experiments, although no OmpW-specific model had been trained, OmpF specificity was predicted for wild-type RB51 but not RB51(Q206L), and for RB68(L206Q) but not wild-type RB68, precisely consistent with the observed phenotypic switch.

Across all engineered variants, predicted and observed phenotypes were in complete agreement, establishing that the sequence features identified by modelling, comparative genomics and structural prediction are the functional determinants of receptor recognition.

## Discussion

This study establishes that phage receptor identity and the sequence determinants of receptor specificity can be inferred from genome sequence once a sufficiently large and systematically phenotyped training set is available. Across 255 *E. coli* phages, genome-wide screens assigned receptor identities to 193 phages, and three orthogonal analyses converged on the molecular basis of those identities. Comparative genomics identified receptor-binding modules and specificity loci, AlphaFold3 localized interface-proximal determinants, and annotation-free *k*-mer models independently recovered the same modules as the dominant predictive features. The experimental scale, encompassing 1,050 genome-wide screens, represents the largest application of barcoded libraries to phage-host specificity to date, exceeding the largest prior single study by more than tenfold. The breadth and diversity of phenotypic data generated here enabled predictive modeling rather than descriptive cataloguing of individual phage-host interactions^24–27,46,47^. Applied to the 1,875 *E. coli* phage genomes currently deposited in the NCBI GenBank database, these models yielded receptor predictions for more than half of all accessions, the large majority of which carried no receptor annotation. In several instances, the annotation-free models outperformed comparative genomics, identifying meaningful sequence polymorphisms at the subdomain level down to individual amino acids without requiring prior biological knowledge.

The systematic screens revealed that only 13 of the 60 to 180 outer membrane proteins estimated in the *E. coli* K-12 genome^48,49^ are recognized by phages, including two receptors not previously associated with phage infection, OmpW and NupG. The restricted size of this functional receptor repertoire likely reflects evolutionary convergence on a specific class of targets, namely OMPs that are broadly conserved across the *E. coli* pangenome^50^, highly expressed, and centrally positioned in the cellular protein interaction network^51^. These properties also characterize receptor proteins of mammalian viruses^52^, suggesting similar selective constraints operate on virus-host recognition across domains of life. That many phage-targeted OMPs perform essential physiological functions further limits the evolutionary options available to bacteria seeking to escape predation through receptor loss or modification^53^.

On the phage side, receptor-recognizing modules are organized in designated specificity loci, conserved within families and frequently exchanged by horizontal gene transfer between distant taxa, with the coexistence of diverse phages on clonal hosts likely facilitating these exchanges and driving diversification^54,55^. The identification of a conserved NGR-binding domain in otherwise unrelated tail proteins of myoviruses and podoviruses further suggests that convergent evolution can independently produce equivalent recognition solutions. The receptor-binding domains identified by comparative genomics and structural modelling were independently recovered as the dominant predictive features by annotation-free sequence models, providing cross-validation across three methodologically distinct approaches. Phage engineering experiments confirmed this directly, with domain swaps between Gp38 adhesins and Gp12 short tail fibers redirecting primary and secondary receptor specificity as predicted, combinatorial exchanges generating receptor combinations absent from all wild-type phages, and bidirectional mutagenesis of Q206L establishing that a single residue change is both necessary and sufficient to switch OmpF specificity to OmpW. This single amino acid substitution demonstrates that phenotypic innovation in phage-host recognition can arise from minimal genetic change, with implications for both understanding receptor evolution in natural populations and designing targeted engineering strategies^56^.

Having demonstrated that systematic phenotyping at scale enables reliable sequence-based prediction of receptor specificity, extending this framework beyond model *E. coli* strains to clinically relevant pathogenic bacteria represents the most immediate priority. O-antigen and capsular polysaccharides, absent from the strains used here except for the stress-induced colanic acid capsule, are major virulence determinants^57^ and phage receptors^58^ in *E. coli* and in Gram-negative ESKAPE pathogens including *Klebsiella pneumoniae*^59^, *Pseudomonas aeruginosa*^60^ and *Acinetobacter baumannii*^61^. These surface structures are extensively diverse across bacterial populations^62–64^ and their recognition is mediated by highly specific receptor-binding proteins whose sequence determinants remain largely uncharacterized^65,66^. In a recent study extending this framework to a wild-type *E. coli* strain expressing OPS and CPS, genome-wide screens identified which phages required polysaccharide recognition for infection and which were sterically blocked from accessing deeper receptors by their presence, demonstrating that the approach is applicable to polysaccharide-expressing hosts^43^. Building equivalent datasets in such hosts and training models on diverse OPS and CPS variants would address one of the most consequential gaps in current phage therapy selection strategies^8,67^. This framework is also complementary to recent work predicting phage-host interactions at the strain level^10^: such approaches predict whether a phage infects a given strain, whereas ours predicts which molecular receptor it targets and which sequence features determine that targeting. Testing our characterized phage collection against a large panel of *E. coli* strains would further clarify the contribution of receptor specificity to host range and inform the engineering of phages targeting defined bacterial subsets. More broadly, the establishment of scalable phenotyping and predictive modelling frameworks will be essential to implementing phage-based solutions across health, environmental biotechnology and microbiome editing.

## Methods

### Bacterial strains, plasmids and growth conditions

All the bacterial strains and plasmids used in this study are listed in **Tables S2** and **S10**, respectively. Unless stated otherwise, bacterial strains were grown in LB broth or on LB agar plates, and incubation was carried out at 37 °C, with liquid cultures being shaken at 200 rpm. When required, antibiotics were added at a final concentration of 25 µg/mL for kanamycin (Kan) and 30 µg/mL for chloramphenicol (Cam), while calcium chloride (CaCl_2_) was supplemented at 10 mM for phage experiments. Bacterial strains were kept for long-term storage at -80 °C in 15 % sterile glycerol. Plasmids and genomic DNA were routinely stored at -20 °C. All enzymes were acquired from New England Biolabs (NEB) while primers were synthesized at Elim Biopharmaceuticals and Integrated DNA Technologies (IDT).

### Phage acquisition and isolation

*Escherichia coli* dsDNA phages were obtained from public and personal collections, or isolated in-house from environmental samples (**Table S1**). Phages reported in previous studies were kindly provided by Prof. Alexander Harms at the Biozentrum, University of Basel, Switzerland^18,19^, Prof. Sylvain Moineau at the Félix d’Hérelle Reference Center for Bacterial Viruses, Université Laval, QC, Canada^68^, Prof. Jennifer Mahony at the School of Microbiology and APC Microbiome Ireland, University College Cork, Ireland^69^, Dr. Marie-Agnès Petit at the Micalis Institute, University Paris-Saclay, INRAE, AgroParisTech, France^70^, Prof. Jeremy J. Barr at the School of Biological Sciences, Monash University, Australia^71^, Dr. Andrew D. Brabban and Dr. Elizabeth Kutter at the Evergreen State College, WA, USA^72–74^, Prof. Petr Leiman at the Department of Biochemistry & Molecular Biology, University of Texas Medical Branch, USA^42^, Dr. Nora Pyenson at the Grossman School of Medicine, New York University, NY, USA^54^, Prof. Ben Temperton at the Faculty of Health and Life Sciences, University of Exeter, UK^75^ and Prof. Andrew Millard at the Department of Genetics and Genome biology, University of Leicester, UK^76^. Jumbo phages Goslar and G17^77^ were acquired from the DSMZ collection (https://www.dsmz.de/collection). Additional phages were also kindly shared by Prof. Stephen T. Abedon from his collection at the Department of Microbiology of the Ohio State University, USA^78^. Classical *E. coli* dsDNA phages (T-phages T2 to T7, P1vir, P2vir, N4 and Lambda) were already available in our lab^24^.

Thirty-one phages were also isolated from diverse environmental sources using several wild-type *E.* coli isolates (**Tables S1** and **S2**). Briefly, phages were enriched by mixing 40 µL of an overnight bacterial culture with 2 mL of 0.22 µm-filtered environmental sample and 2 mL of 2X LB supplemented with CaCl_2_ into a 24 deep well plate sealed with a Breathe-Easy® membrane (Diversified Biotech) and incubating overnight under agitation at 37 °C. Enrichments were then centrifuged at 2,000 x g for 10 min and 100 µL of the supernatant was filtered on 0.22 µm with an AccroPrep™ 96-well filter plate (Cytiva) by centrifuging again. The resulting filtrates were mixed with 100 µL on an overnight culture of the host into 5 mL of 0.5 % Tryptone top agar (T-top) supplemented with CaCl_2_, poured onto agar plates and incubated overnight. Phage plaques were then subjected to three rounds of streak-purification before being amplified into large volumes.

### Phage amplification and titration

Phage amplification was carried out from single phage plaques cultivated with their respective hosts (**Table S1**). An overnight culture of the bacterial host was diluted to an OD_600_ of 0.1 in 35 mL of LB broth supplemented with CaCl_2_ in a 125-mL flask. Isolated plaques were inoculated and the flasks were incubated under agitation until complete clearance was observed, or up to 4 h otherwise. Immediately after, 100 µL of chloroform (CHCl_3_) was added to the flask and lysates were incubated at room temperature for 10 min. Lysates were transferred to 50-mL Falcon tubes and centrifuged for 10 min at 20,913 x g to pellet cell debris and chloroform. The supernatants were then filtered on 0.22-µm Millex® polyethersulfone syringe filters (Millipore-Sigma) and stored at 4 °C. Determination of phage titers was carried out in high-throughput by serially-diluting up to 12 lysates to 10^-8^ in a 96-well PCR plate in a final volume of 200 µL of SM buffer (Teknova) supplemented with CaCl_2_. A bacterial lawn of the relevant host strain was prepared from an overnight culture following the standard double-layer agar method using 0.5 % T-top agar supplemented with CaCl_2_ in a rectangular Nunc™ Omnitray™ single-well plate (Thermo Scientific). Using a Rainin MicroPro 300 semi-automated pipettor (Mettler-Toledo Rainin), 2 µL droplets of phage dilutions were spotted in triplicates onto the bacterial lawn, allowed to dry and plates were incubated overnight. Plates were scanned the next morning to enumerate phage plaques over the triplicates and calculate the titers.

### Genome sequencing, assembly and annotation

Whole-genome sequencing was conducted on all phages to generate new genome sequences when necessary, and to check the identity and purity of our phage stocks as part of our quality control process. DNA extractions were performed from 1 mL of phage lysate using the Phage DNA Isolation Kit (Norgen Biotek Corp). Briefly, remaining bacterial nucleic acids were removed prior to the extraction by incubating the lysate at 37 °C for 30 min with a nuclease mix containing 2 µL of RNase A/T1 mix (2 mg/mL and 5000 U/mL, respectively), 2 µL of DNAse I (1 U/µL) and 1 µL of 10X DNAse buffer (Thermo Scientific). Nucleases were then inactivated by incubating at 80 °C for 5 min. Phage lysis was carried out by mixing the suspension with 500 µL of the kit’s lysis buffer B and 4 µL of proteinase K at 20 mg/mL (QIAgen), followed by incubating 15 min at 55 °C and another 15 min at 65 °C. Subsequent steps were conducted following the manufacturer’s protocol. Phage genomes were sequenced on Illumina MiSeq platforms with a target coverage of 15 – 30 X at Neochromosome (New York, USA) and NGS Lab (New York, USA) in 300-bp paired-end, and at QB3 Genomics (UC Berkeley, Berkeley, CA, RRID:SCR_022170) in 150-bp paired-end. Raw reads were trimmed with trim_galore v0.6.10^79^ with default parameters, and bbduk v39.06-0^80^ (options “qtrim=r trimq=20 maq=20”). Next, reads were normalized to a target coverage of 100x (bbnorm v39.06-0, target=100 min=2) and assembled with spades v4.2.0^81^ (options “-- isolate” and “-k 21, 33, 55, 77, 99, 121”). From the assembly, contigs of ≥ 2 kb were screened with geNomad v.1.9.4^82^ to distinguish phage from host contigs (options “end-to-end” and “--conservative”), and predicted phage contigs were further checked for quality and host contamination wich CheckV v1.0.3^83^ (options “end-to-end”). For each library, the contig identified as representing the complete phage genome was then selected for annotation. Phage genome assemblies were annotated with Pharokka v1.6.1^84^. Specifically, coding sequences (CDS) were predicted with PHANOTATE^85^, tRNAs were predicted with tRNAscan-SE 2.0^86^, tmRNAs were predicted with Aragorn^87^ and CRISPRs were predicted with CRT^88^. Functional annotation was generated by matching each CDS to the PHROGs^89^, VFDB^90^ and CARD^91^ databases using MMseqs2^92^ and PyHMMER^93^. Contigs were matched to their closest hit in the INPHARED database^94^ using mash^95^. Annotations of coding genes were improved with PHOLD v 0.2.0^96^. Available NCBI assemblies were sometimes preferred over newly-generated sequences if the quality or completeness was better, but all assemblies were systematically re-annotated with the same pipeline. Proteomic equivalence quotient (PEQ) between phage pairs was calculated with PhamClust v1.3.3^30^. Phage gene-sharing networks were computed with vConTACT 2.0^29^, and annotated and visualized with Cytoscape v.3.10.4^97^.

Bacterial DNA was extracted from 1 mL of overnight culture using the Wizard® Genomic DNA Purification Kit (Promega Corporation) according to the manufacturer’s protocol. Bacterial genome sequencing was then carried out by Plasmidsaurus using Oxford Nanopore Technology with custom analysis and annotation.

### Genome-wide competitive phage fitness assays

RB-TnSeq libraries previously built in *E. coli* BW25113 and BL21 and a Dub-seq library constructed in BW25113 were used to massively screen for bacterial genes associated with phage resistance in liquid cultures^24^. First, a 1-mL aliquot of mutant library was thawed and inoculated into 25 mL of LB supplemented with kanamycin (for RB-TnSeq) or chloramphenicol (for Dub-seq), and grown under agitation at 37 °C until reaching an OD_600_ of 3 – 4. Three 1-mL samples were immediately taken and centrifuged for 5 min at 15,900 x g to collect cell pellets to serve as reference for the BarSeq (labelled as time zero), and immediately stored at -20 °C. The recovered mutant library was then diluted in 2X LB to reach an OD_600_ of 0.04, corresponding to ∼ 4 x 10^7^ CFU/mL. Chloramphenicol was also added in 2X LB for the Dub-seq experiments. Sterile 48-well tissue culture plates were then filled by mixing in each well 350 µL of the diluted mutant library and 350 µL of phage suspensions normalized at ∼10^9^ PFU/mL in sterile SM buffer supplemented with CaCl_2_ in order to achieve a multiplicity over infection (MOI) range of 10 – 100. Controls without phage were also performed by mixing the mutant library with 350 µL of phage buffer. The plates were then sealed with a Breathe-Easy® sealing membrane and incubated at 37 °C with orbital shaking for 14h – 16h in a BioTek 800 TS plate reader (Agilent Technologies), with OD_630_ readings every 15 min. Surviving cells were collected by centrifugation and immediately processed for DNA or plasmid extraction, or stored for later use at -20 °C. All phage assays were at minimum conducted in duplicate.

### BarSeq sample preparation, sequencing and data processing

Genomic DNA extractions of RB-TnSeq cell pellets were performed in high-throughput using the QIAamp 96 DNA QIAcube HT Kit (QIAgen) following the manufacturer’s protocol. RNase treatment was carried out by adding 20 µL of RNaseA at 20 mg/mL (Sigma-Aldritch) to cell lysates and incubating at 37 °C for up to 1 h. Elution was performed in a 96-Elution CL Microtubes plate (QIAgen) with 100 µL of AE elution buffer and 10 µL of TopElute buffer. For Dub-seq experiments, high-throughput plasmid minipreps were similarly conducted with the QIAprep 96 Plus Miniprep Kit (QIAgen). BarSeq PCRs and Illumina sequencing were carried out following established procedures^24^. Bacterial mutants were considered to be significantly enriched and having a high fitness after phage exposure when the log_2_ fold-change (log_2_FC) of read abundance was above or equal to 4, and significantly depleted causing a low fitness when the log_2_FC was below or equal to -2. In both cases, statistical significance was established when the t-statistic absolute value was above or equal to 5^22^. Fitness data of individual transposon mutants was viewed in an internal instance of the Fitness Browser^98^, and with the dedicated BarSeq browser presented in this study. Graphic visualization of the genome-wide screens results was achieved with the ggplot2 v4.0.0^99^ and ggtree v3.10.1^100^ R packages.

### Phenotypic validations

In order to validate the contribution of some *E. coli* BW25113 genes to phage susceptibility deduced from RB-TnSeq screens, single-gene deletion mutants from the KEIO collection^31^ containing the pCA24N empty vector (as a control) or corresponding ASKA IPTG-inducible complementation plasmid^32^ were used to test the susceptibility to subsets of phages. Similarly, confirmation of Dub-seq hits were conducted directly using the ASKA collection, as these strains’ genetic background is the same as our Dub-seq library’s. All the ASKA plasmids were extracted using the QIAprep Spin Miniprep Kit (QIAgen), and plasmids were chemically-transformed into KEIO strains following standard cloning protocol^101^. Transformant colonies bearing the plasmids were selected on LB agar plates supplemented with kanamycin and chloramphenicol. For subsequent phage susceptibility assays, the bottom LB agar was supplemented with kanamycin, chloramphenicol, and 0 – 0.1 mM IPTG. Subsets of phages were serially-diluted and spotted following the same protocol used for phage titration (see **Phage amplification and titration**). Phage susceptibility tests were performed in technical duplicates, and the efficiency of plating (EOP) was calculated as the mean of the ratio of the estimated number of single plaques on the deletion mutant over the number of plaques observed on the parental strain. Restoration of the parental phenotype when complemented with ASKA plasmids was systematically checked as a control.

### Phage receptor-binding proteins and domains identification

If not known from prior evidence, putative receptor-binding protein (RBP) genes were manually identified in phage genomes by searching for gene(s) presence/absence matching the observed adsorption phenotypes. Protein homology with known RBPs was searched with PaperBLAST^102^ and homology between putative RBPs was determined against an internal phage genome database with BLASTp^103^. Phage genome synteny plots were generated with the gggenomes v1.1.3^104^ R package. Multiple protein sequence alignments were generated in Seaview v5.1^105^ with Clustal Omega v1.1.0^106^ and visualized with the ggmsa v1.0.3^107^ R package. Binding of putative RBPs to host outer membrane proteins was modeled with AlphaFold3^6^ on the AlphaFold Server (https://alphafoldserver.com), and protein-protein models were then visualized and annotated with ChimeraX v1.11.1^108^. When multiple putative RBPs were considered, the protein pair showing the highest ipTM value and appropriate directionality was determined to be the *bona fide* true RBP.

### Receptor-binding protein swapping

Selected phage RBPs were fully or partially exchanged by homologous recombination between genetically similar phage isolates to validate their causal roles in the adsorption phenotypes. All constructs designed for homologous recombination were synthesized *de novo* and cloned into the pTwist Chlor High Copy vector at Twist Biosciences (**Table S10**). All plasmids were transformed into BW25113 following standard cloning protocol^101^, and selection of recombinant phages was achieved using relevant KEIO single-gene deletion mutants^31^. A first design (pLM1) consisted of the complete *gp38* adhesin gene of phage EV240 flanked by the upstream and downstream 250 bp adjacent regions of phage CM8’s *gp38* to obtain a CM8×EV240 recombinant binding OmpA (selection on BW25113Δ*tsx*). A second construct (pLM2) was designed to have the *gp12* tailspike gene of phage JK38 encompassed between the upstream and downstream 250 bp adjacent regions of CM8’s *gp12* to generate a CM8×JK38 recombinant binding the Kdo sugar of the LPS inner core (selection on BW25113Δ*waaC*). The CM8 double mutant containing EV240’s *gp38* and JK38’s *gp12* was generated from the CM8×JK38 recombinant phage using the same protocol used to obtain the CM8×EV240 recombinant mutant. Similarly, two additional constructs were designed to confirm the role of mutations in the unique adsorption phenotypes of phage RB68 compared to its clonal relative RB51. First, a 301 bp fragment of RB51’s *gp38* centred around the adenine in position 617 of the gene was cloned in construct pLM5 in order to obtain a RB68×RB51 recombinant binding OmpF (selection on BW25113Δ*ompW*). Conversely, the same 301 bp fragment of RB68’s gp38 which contains a thymine in position 617 was built into construct pLM6 to generate a RB51×RB68 recombinant binding OmpW (selection on BW25113Δ*ompR*).

Selection of recombinant phages was carried out by infecting pLM plasmids-containing BW25113 cultures with the corresponding phages and selecting on single-gene deletion mutants. Briefly, 100 µL of an overnight bacterial culture grown in LB supplemented with chloramphenicol was mixed in 5 mL of T-top agar with 100 µL of the relevant phage diluted at 10^4^ PFU/mL (to obtain ∼ 1000 PFU). The suspension was then poured onto LB chloramphenicol plates and incubated overnight. Negative controls were systematically performed by infecting a plasmid-free BW25113 culture in parallel. The next day, plates were flooded with 5 mL of SM buffer supplemented with CaCl_2_ and scrapped to detach and break down the top agar layer. The resulting slurry was transferred to a 15-mL Falcon tube, vortexed thoroughly to mechanically release phages, centrifuged for 5 min at 20,913 x g and the supernatant was diluted to 10^-2^ in buffer. One hundred µL of the pure and diluted lysates were then similarly mixed in 5 mL T-top agar with 100 µL of the appropriate single-gene deletion mutant overnight cultures grown in LB supplemented with kanamycin, poured onto LB kanamycin plates and incubated overnight. Lysates produced with the plasmid-free BW25113 were also tested on the mutants to control for the absence of plaques. Recombinant phage plaques were picked the next day and subjected to two additional rounds of streak-purification on their corresponding mutants to ensure their purity and absence of remaining wild-type phages. Recombinant phages were then amplified as described above (see **Phage amplification and titration**), and their whole genome was sequenced to confirm the mutations and verify the absence of secondary mutations in the genomes (see **Genome sequencing, assembly and annotation**). Finally, the new adsorption specificity phenotypes were confirmed by RB-TnSeq and by determining their EOP on relevant KEIO mutants as described in **Genome-wide competitive phage fitness assays** and **Phenotypic validations**.

### Host target predictive modelling

#### Predictive modeling

Phage receptor prediction was performed using GenoPHI v0.1, a package developed for genotype-to-phenotype prediction of phage-host interactions^43^. GenoPHI represents phage proteomes as presence-absence matrices of amino acid *k*-mer features, enabling identification of predictive *k*-mer profiles through recursive feature elimination (RFE) and downstream classification. Each phage was assigned one or several receptors, including OMPs, the NGR, and LPS core sugars, based on RB-TnSeq data. This resulted in varying class imbalance across receptors, from 41 and 40 phages binding BtuB and Kdo, respectively, to 6 and 5 phages binding LamB and HepII, respectively. Receptors targeted by fewer than 4 phages were excluded due to insufficient positive-class representation.

We framed receptor prediction as 13 independent binary classification tasks, with each model predicting whether a phage uses a specific receptor (i.e., the outer membrane proteins Tsx, OmpA, OmpF, OmpC, FhuA, BtuB, LptD, LamB; the LPS core sugars Kdo, HepI, HepII, GluI; the NGR polysaccharide). This approach addressed the substantial class imbalance that would confound a single multilabel classifier. Critically, amino acid sequences from all phage genes were used as input, not just known receptor-binding proteins, making the models agnostic to known interaction mechanisms. These sequences were processed through the GenoPHI *k*-mer workflow using *k* = 5 to generate receptor-specific predictive models.

#### Computational cross-validation

Model performance was evaluated independently for each receptor using 20-fold cross-validation, with a random 10% of phages representing both positive and negative classes held out of feature generation, selection, and modeling workflows. Held-out phages were then assigned *k*-mer features and used for prediction. Performance was assessed using Matthews Correlation Coefficient (MCC), accuracy, precision, and recall at a classification threshold of 0.5, as well as area under the receiver operating characteristic curve (AUROC) as a threshold-independent metric.

#### Target prediction on novel phages

Initial models were trained on a subset of 206 phages, and applied to predict receptor specificity in 18,398 phage genomes from the NCBI GenBank database (accessed May 2025), including 49 phage genomes which were selected for experimental cross-validation. Final models were trained on all available phages for each receptor. In all cases, phage proteomes were assigned *k*-mer features using the default GenoPHI workflow, and receptors were predicted across all models using a classification threshold of 0.5.

### Interactive Phage Datasheets, BarSeq browser and Phage Genome Browser

We built the interactive Phage Datasheets homepage as a static site using javascript, d3, cgview for genbank file parsing, and Claude (used primarily for code review and assistance with layout and user interaction). We built the interactive BarSeq browser using javascript, d3, and arquero. For both browsers, data was reformatted from the raw data underlying the study for easy and efficient loading. All code is available at https://github.com/mjohnson11/PhageDataSheets/tree/main/Ecoli_phages, and the browser is available online at https://iseq.lbl.gov/PhageDataSheets/Ecoli_phages/. We built the interactive Phage Genome Browser based on the GenomeDepot platform for genome management and comparative analysis^109^. The browser contains all phage genomes used in this study and is available at https://iseq.lbl.gov/ecoliphages.

## Acknowledgments

The authors thank members of the BRaVE Phage Foundry team, the Arkin lab, the Mutalik lab, Benjamin A. Adler and Morgan N. Price for feedback and discussion, and all of our phage friends who shared phages for this work.

## Author Contributions

**L.M.**: Conceptualization, Methodology, Investigation, Validation, Formal Analysis, Data Curation, Visualization, Writing - Original Draft, Review & Editing

**A.J.C.N.**: Conceptualization, Methodology, Investigation, Software, Formal Analysis, Data Curation, Visualization, Writing - Review & Editing

**A.K.**: Investigation, Software, Formal Analysis, Data Curation, Visualization, Writing - Review & Editing

**M.P.**: Investigation, Validation

**M.S.**: Investigation, Validation

**E.O.R.**: Investigation, Validation

**F.M.**: Investigation, Validation, Formal Analysis, Data Curation, Writing - Review & Editing

**M.S.J.**: Software, Data Curation, Visualization, Writing - Review & Editing

**S. R.**: Investigation, Software, Data Curation, Writing - Review & Editing

**B.K.**: Writing - Review & Editing

**A.M.D.**: Validation, Writing - Review & Editing, Supervision

**E.G.D.**: Writing - Review & Editing, Supervision, Funding Acquisition, Project Administration

**V.K.M.**: Conceptualization, Methodology, Writing - Review & Editing, Supervision, Funding Acquisition, Project Administration

**A.P.A.**: Conceptualization, Methodology, Writing - Review & Editing, Supervision, Funding Acquisition, Project Administration

## Data Availability

Assembled genome sequences for all 260 phages characterized in this study have been deposited in NCBI GenBank under accession numbers [*submitted*]. Bacterial genome sequences are available under accession numbers [*submitted*]. Genome-wide fitness data from all 1,050 RB-TnSeq and Dub-seq screens are available through the BarSeq browser, and filtered high-scoring genes are listed in **Supplementary Datasets S1–S3**. All phage metadata, including receptor assignments, receptor-binding protein annotations, measured EOP values, and AlphaFold3 model metadata are available on the interactive Phage Datasheets and provided in **Supplementary Tables S1** and **S5–S7**, and **Supplementary Datasets S4-S5**. *K*-mer feature sets identified by predictive models and receptor predictions for NCBI phage genomes are provided in **Supplementary Datasets S6–S7**. All supplementary materials (**Supplementary Figures S1-S13**, **Supplementary Tables S1-S10**, **Supplementary Datasets S1-S7**), and high-resolution image files of Figures 1-4 are available on Figshare at https://doi.org/10.6084/m9.figshare.31930314.

## Code Availability

The GenoPHI software package used for *k*-mer-based receptor prediction is available at https://github.com/Noonanav/GenoPHI. Code for the interactive Phage Datasheets and BarSeq browser is available at https://github.com/mjohnson11/PhageDataSheets/tree/main/Ecoli_phages, and the browsers are accessible at https://iseq.lbl.gov/PhageDataSheets/Ecoli_phages/. Code for the Phage Genome Browser is available at https://github.com/aekazakov/genome-depot, and the browser is accessible at https://iseq.lbl.gov/ecoliphages.

## Materials & Correspondence

All other data supporting the findings of this study are available from the corresponding author Vivek K. Mutalik upon reasonable request.

## Supplementary Information

**Figure S1. PEQ heatmap between *E. coli* phages.** Pairwise proteomic equivalence quotient (PEQ) values between the 255 *E. coli* dsDNA phages presented in this study.

**Figure S2. EOP estimation of *Straboviridae* phages on LPS core deletion mutants.** Efficiency of plating (EOP) values of 52 *Straboviridae* phages tested on the LPS core deletion mutants BW25113Δ*waaC* (HepI), BW25113Δ*waaF* (HepII), and BW25113Δ*waaG* (GluI), harboring the empty plasmid pCA24N (blue bars) or expressing the complemented gene *in trans* on the same plasmid (yellow bars). Values are the mean EOP calculated over biological duplicates. Phage name colors correspond to the primary porin receptor targeted by each phage, and the experimentally-characterized secondary receptor (Kdo or HepI) and phage genera are indicated above.

**Figure S3. Experimental validation of the dual-receptor phenotypes exhibited by the *Drexlerviridae* phages Bas01 and CPL00221.** (**A**) Efficiency of plating (EOP) values of Bas01 and CPL00221 on the outer membrane protein (OMP) deletion mutants BW25113Δ*nupG* and BW25113Δ*fhuA*, and the spontaneous phage-resistant mutant BW25113*lptD*Δ(Y658-Y678)::Y. Each mutant is harboring the empty plasmid pCA24N or expressing the complemented gene *in trans* on the same plasmid. Values are the mean EOP calculated over biological duplicates. It should be noted that complementation of *lptD* failed to restore the wild-type phenotype, putatively because expressing the gene *in trans* does not effectively replace the native truncated LptD proteins expressed by the cell. (**B**) Pictures of the plaques obtained when spotting 10-fold dilutions of Bas01 and CPL00221 on the 3 OMP mutants, with or without complementation *in trans*.

**Figure S4. RB-TnSeq-based receptor identification in other myoviruses.** Annotated heatmap showing the fitness values (log_2_ fold-change) for 34 high-scoring genes, the related membrane structures, and the resulting receptor(s) assignment in 56 non-*Straboviridae* myoviruses. Phages included in the model testing subset are highlighted in red. Grey boxes indicate missing data.

**Figure S5. RB-TnSeq-based receptor identification in podoviruses.** Annotated heatmap showing the fitness values (log_2_ fold-change) for 23 high-scoring genes, the related membrane structures, and the resulting receptor(s) assignment in 25 podoviruses. Grey boxes indicate missing data.

**Figure S6. TolC- and LamB-specific GpJ C-terminal domains identified by comparative genomics.** Protein alignment of the C-terminal domains of GpJ for 2 TolC- and 5 LamB-specific *Drexlerviridae* and related phages (alignment positions 1388 to 1572 / 1572 total). The central tail fiber GpJ of Lambda phage was not included as its C-terminal domain shared little to no homology with the 7 phages displayed. Relevant taxonomic levels and targeted receptors are indicated along the phage names. Amino acid residues are colored according to the “Zappo” color scheme.

**Figure S7. NupG-specific GpJ C-terminal domains identified by comparative genomics.** Protein alignment of the central tail fiber GpJ C-terminal domains of 21 FhuA-, 18 BtuB-, 12 LptD-, 4 YncD-, and 2 NupG-specific *Drexlerviridae* and related phages (alignment positions 885 to 1146 / 1329 total). Relevant taxonomic levels and targeted receptors are indicated along the phage names. Predictive *k*-mer features identified by the FhuA binary classifier are highlighted by red boxes, and amino acid sequences identified by comparative genomics to be unique to the two NupG-specific phages (Bas01 and CPL00221) are highlighted by yellow boxes with dashed red outlines. Amino acid residues are colored according to the “Zappo” color scheme.

**Figure S8. NGR-specific homologous tail fiber domain identified by comparative genomics and annotation-free modelling.** Protein alignment of the putative NGR-binding homologous domain identified in the tail fiber genes of 27 myoviruses and 2 podoviruses (alignment positions 409 to 503 / 1504 total). Alignment is viewed in the “consensus = true” mode, i.e. every amino acid residue varying from the majority consensus is displayed in a white box with grey letters. Relevant taxonomic levels and morphotype information are indicated along the phage names. The predictive and perfectly conserved *k*-mer feature ‘GMSHY’ identified by the NGR binary classifier is highlighted with a black outline. Amino acid residues are colored according to the “Zappo” color scheme.

**Figure S9. ROC curves for the LPS core sugar binary classifiers.** Receiver operating characteristic (ROC) curves showing the sensitivity (true positive rate) and specificity (true negative rate) of the 4 LPS core sugar-specific binary classifiers (Kdo, HepI, HepII, GluI).

**Figure S10. OmpC-specific Gp37 C-terminal domains identified by annotation-free modelling.** Protein alignment of the Gp37 C-terminal domains of the 17 OmpC-specific *Straboviridae* phages (alignment positions 865 to 1118 / 1118 total). Alignment is viewed in the “consensus = false” mode, i.e. every amino acid residue varying from the majority consensus is displayed in colors. The structural domains D10 and D11, and the D11 “stem” and “tip” sub-domains described by Bartual *et al.* 2007 and Islam *et al.* 2019 are indicated above the alignment track. Contiguous predictive *k*-mers identified by the OmpC binary classifier are highlighted by red boxes for phages T4, Bas40, and TP6. Amino acid residues are colored according to the “Zappo” color scheme.

**Figure S11. AlphaFold3 models of representative instances of the Kdo- (JK38) and HepI-binding (CM8) Gp12 short tail fibers of *Straboviridae* phages.** Each model consists of a trimer of Gp12, and monomer chains are indicated by different colors. Contiguous predictive *k*-mers identified by the Kdo and HepI binary classifiers are highlighted on the models in yellow. The C-terminal domains at the bottom of the models are putatively interacting with the targeted LPS core sugars, while the N-terminal domains at the top are connected to the phage baseplate.

**Figure S12. BtuB-, FhuA- and LptD-specific GpJ C-terminal domains identified by annotation-free modelling.** Protein alignment of the central tail fiber GpJ C-terminal domains of 21 FhuA-, 18 BtuB-, 12 LptD-, 4 YncD-, and 2 NupG-specific *Drexlerviridae* and related phages (alignment positions 1256 to 1329 / 1329 total). Relevant taxonomic levels and targeted receptors are indicated along the phage names. Contiguous predictive *k*-mer features identified by the BtuB, FhuA, and LptD binary classifiers are highlighted by red, yellow, and blue boxes, respectively. Amino acid residues are colored according to the “Zappo” color scheme.

**Figure S13. LPS core sugar specificity predictions across NCBI *E. coli* phages.** vConTACT2 gene-sharing network displaying predictions of the 4 LPS core sugars binary classifiers (Kdo, HepI, HepII, and GluI) across 1,875 *E. coli* dsDNA phages from the NCBI GenBank database and the 255 phages in our collection. Phages included in the initial training set (206 phages) are represented with a circle, while phages selected for the testing subset (49 phages) are indicated by a diamond. Colors indicate the predicted receptor for NCBI phages or experimentally determined receptor for phages in the training and testing subsets.

**Table S1. Phages used in this study.** List of the 260 phages (255 wild-types and 5 mutants) used in this study, and their associated metadata.

**Table S2. *Escherichia coli s*trains used in this study.** List of the *E. coli* strains used as hosts for phage isolation or amplification, and the KEIO single-gene deletion mutants.

**Table S3. Computational cross-validation metrics of the 13 binary classifiers trained for receptor prediction.** Number of positive instances is indicated for each receptor class.

**Table S4. Mutations in phage RB68 compared to RB51.** List of all the genetic polymorphisms found when comparing the genome of the OmpC-specific phage RB68 to its OmpF-specific counterpart RB51.

**Table S5. AlphaFold3 confidence values and directionality assessment for 163 OMP-RBP models.** Model directionality was assessed by verifying that the C-terminal domains of the putative RBP were interacting with the outer membrane-exposed domains of the host outer membrane protein receptor.

**Table S6. AlphaFold3 ipTM ranges for 163 OMP-RBP models, sorted by phage taxa and directionality.** The total number of tested OMP-RBP pairs, and the numbers of sound and incoherent OMP-RBP pairs are indicated for each group.

**Table S7. AlphaFold3 confidence values and directionality assessment for 33 non-cognate OMP-RBP control pairs.** Negative control models were tested by modelling known non-cognate OMP-RBP pairs in cohesive phage taxa (e.g., *Straboviridae* RBPs were only tested against all *Straboviridae* potential host receptors).

**Table S8. Model predictions and experimental validations in testing phage subset (49 phages).** Model predictions and experimental receptor assessment are indicated for each of the 49 phages in the subset. Predictions were deemed accurate if no prediction was made and (i) there was no receptor to predict, or (ii) no binary classifier was trained for this specific receptor.

**Table S9. Model predictions and experimental receptor characterization in the engineered phages.** Model predictions and experimental receptor assessment are indicated for each of the 5 wild-types and 5 mutant phages. Predictions were deemed accurate if no prediction was made and (i) there was no receptor to predict, or (ii) no binary classifier was trained for this specific receptor.

**Table S10. Plasmids used in this study.** List of all the plasmids used in this study, whether for complementation of single-gene deletion mutants (ASKA plasmids) or generation of recombinant phages with through targeted RBP swapping.

**Dataset S1. High-scoring genes in BW25113 RB-TnSeq phage experiments.** List of all the BW25113 genes showing significant high fitness scores (fitness ≥ 4; t-test ≥ 5; read count in experiment ≥ 500) in RB-TnSeq phage experiments.

**Dataset S2. High-scoring genes in BW25113 Dub-seq phage experiments.** List of all the BW25113 genes showing significant high fitness scores (fitness ≥ 4) in Dub-seq phage experiments.

**Dataset S3. High-scoring genes in BL21 RB-TnSeq phage experiments.** List of all the BL21 genes showing significant high fitness scores (fitness ≥ 4; t-test ≥ 5; read count in experiment ≥ 500) in RB-TnSeq phage experiments.

**Dataset S4. EOP values on KEIO single gene deletion mutants.** List of all the efficiency of plating (EOP) values (mean over biological duplicates) obtained by spotting dilutions of individual phages on KEIO single-gene deletion mutants with or without complementation by an ASKA plasmid.

**Dataset S5. EOP values on ASKA single gene overexpression strains.** List of all the efficiency of plating (EOP) values (mean over biological duplicates) obtained by spotting dilutions of individual phages on ASKA single-gene overexpression mutants.

**Dataset S6. Lists of predictive k-mer features identified by the 13 binary classifiers.** Complete list of the predictive *k*-mer features identified by the 8 OMP-, the 4 LPS core sugar- and the NGR-specific binary classifiers across the 255 phages used in this study.

**Dataset S7. Receptor predictions across 18,398 NCBI Caudoviricetes phages.** Summary of the receptor predictions made by the 13 binary classifiers across every tailed phage genome retrieved from the NCBI GenBank database.

