## Supplementary Figures S1-S13 for "Enabling the prediction of phage receptor specificity from genome data"

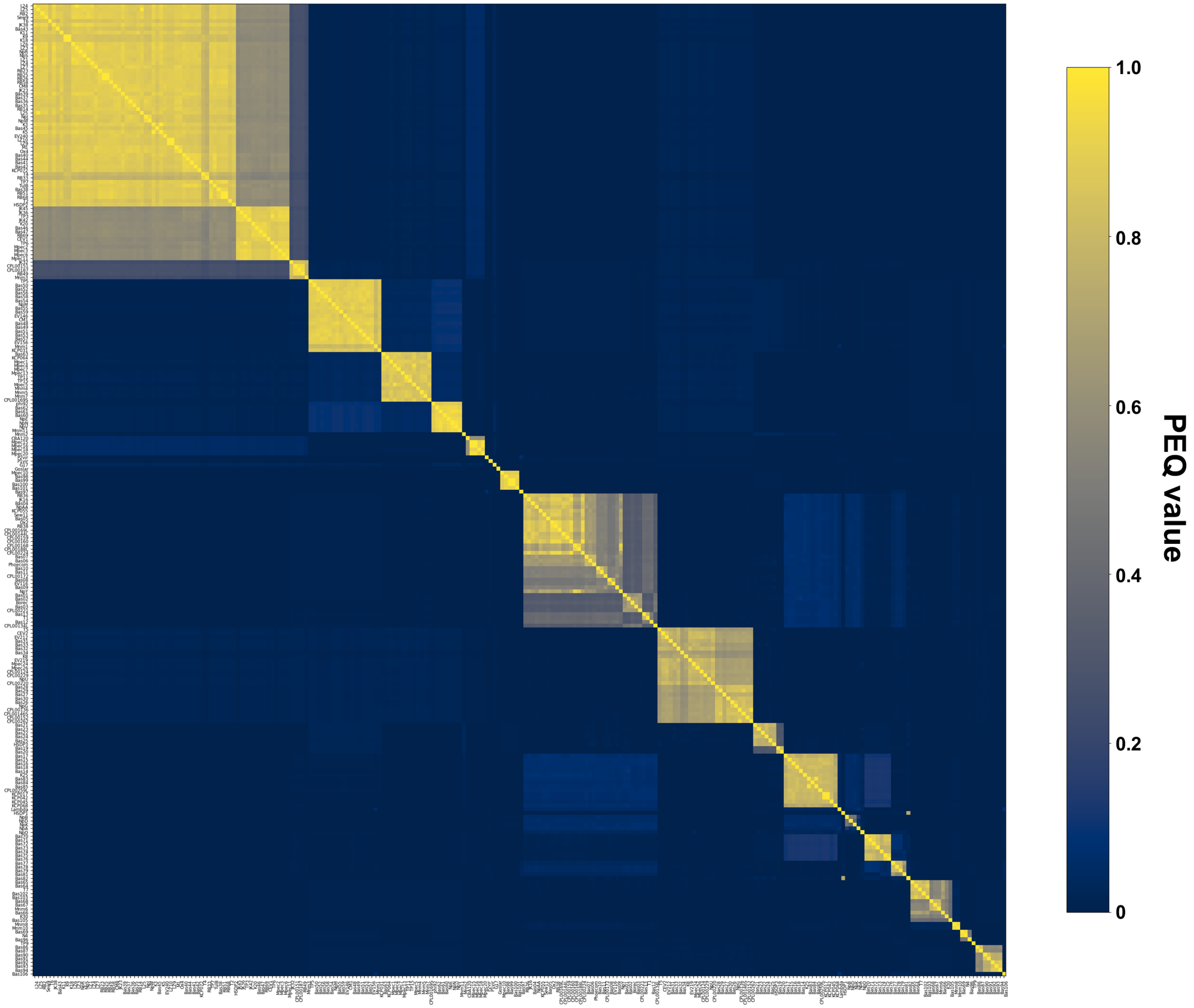


**Figure S1. PEQ heatmap between *E. coli* phages.** Pairwise proteomic equivalence quotient (PEQ) values between the 255 *E. coli* dsDNA phages presented in this study.

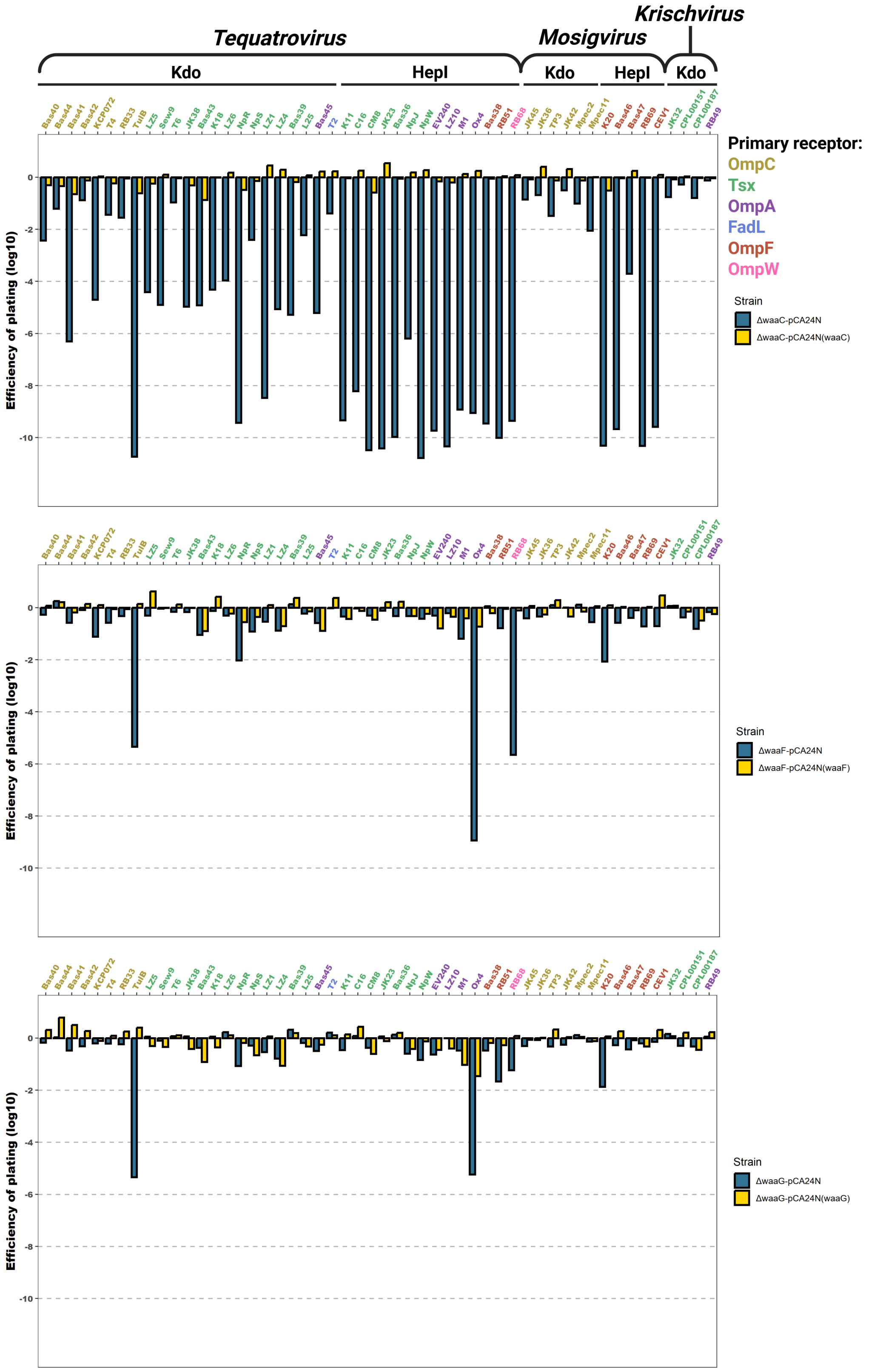


**Figure S2. EOP estimation of *Straboviridae* phages on LPS core deletion mutants.** Efficiency of plating (EOP) values of 52 *Straboviridae* phages tested on the LPS core deletion mutants BW25113Δ*waaC* (HepI), BW25113Δ*waaF* (HepII), and BW25113Δ*waaG* (GluI), harboring the empty plasmid pCA24N (blue bars) or expressing the complemented gene *in trans* on the same plasmid (yellow bars). Values are the mean EOP calculated over biological duplicates. Phage name colors correspond to the primary porin receptor targeted by each phage, and the experimentally-characterized secondary receptor (Kdo or HepI) and phage genera are indicated above.

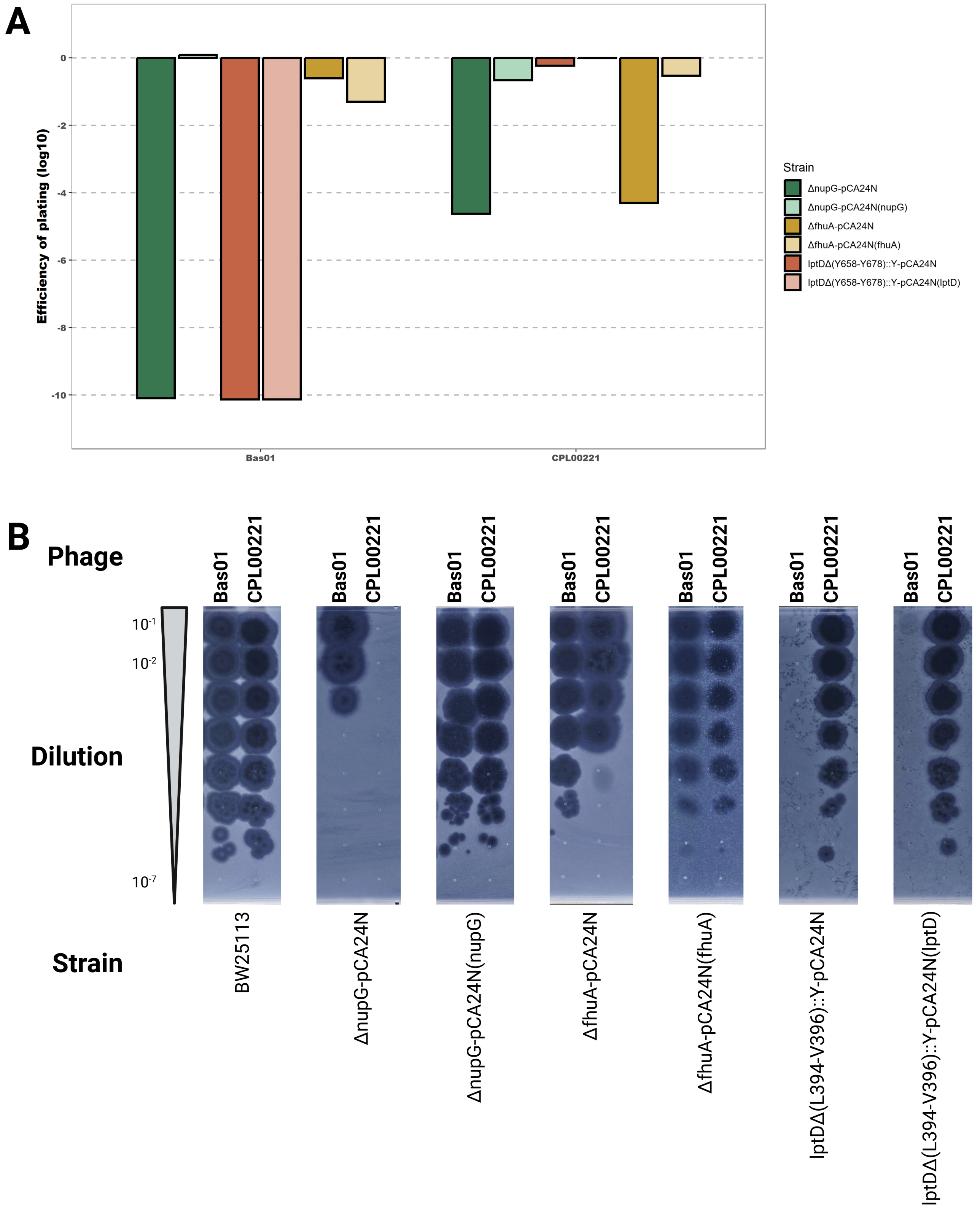


**Figure S3. Experimental validation of the dual-receptor phenotypes exhibited by the *Drexlerviridae* phages Bas01 and CPL00221.** (**A**) Efficiency of plating (EOP) values of Bas01 and CPL00221 on the outer membrane protein (OMP) deletion mutants BW25113Δ*nupG* and BW25113Δ*fhuA*, and the spontaneous phage-resistant mutant BW25113*lptD*Δ(Y658-Y678)::Y. Each mutant is harboring the empty plasmid pCA24N or expressing the complemented gene *in trans* on the same plasmid. Values are the mean EOP calculated over biological duplicates. It should be noted that complementation of *lptD* failed to restore the wild-type phenotype, putatively because expressing the gene *in trans* does not effectively replace the native truncated LptD proteins expressed by the cell. (**B**) Pictures of the plaques obtained when spotting 10-fold dilutions of Bas01 and CPL00221 on the 3 OMP mutants, with or without complementation *in trans*.

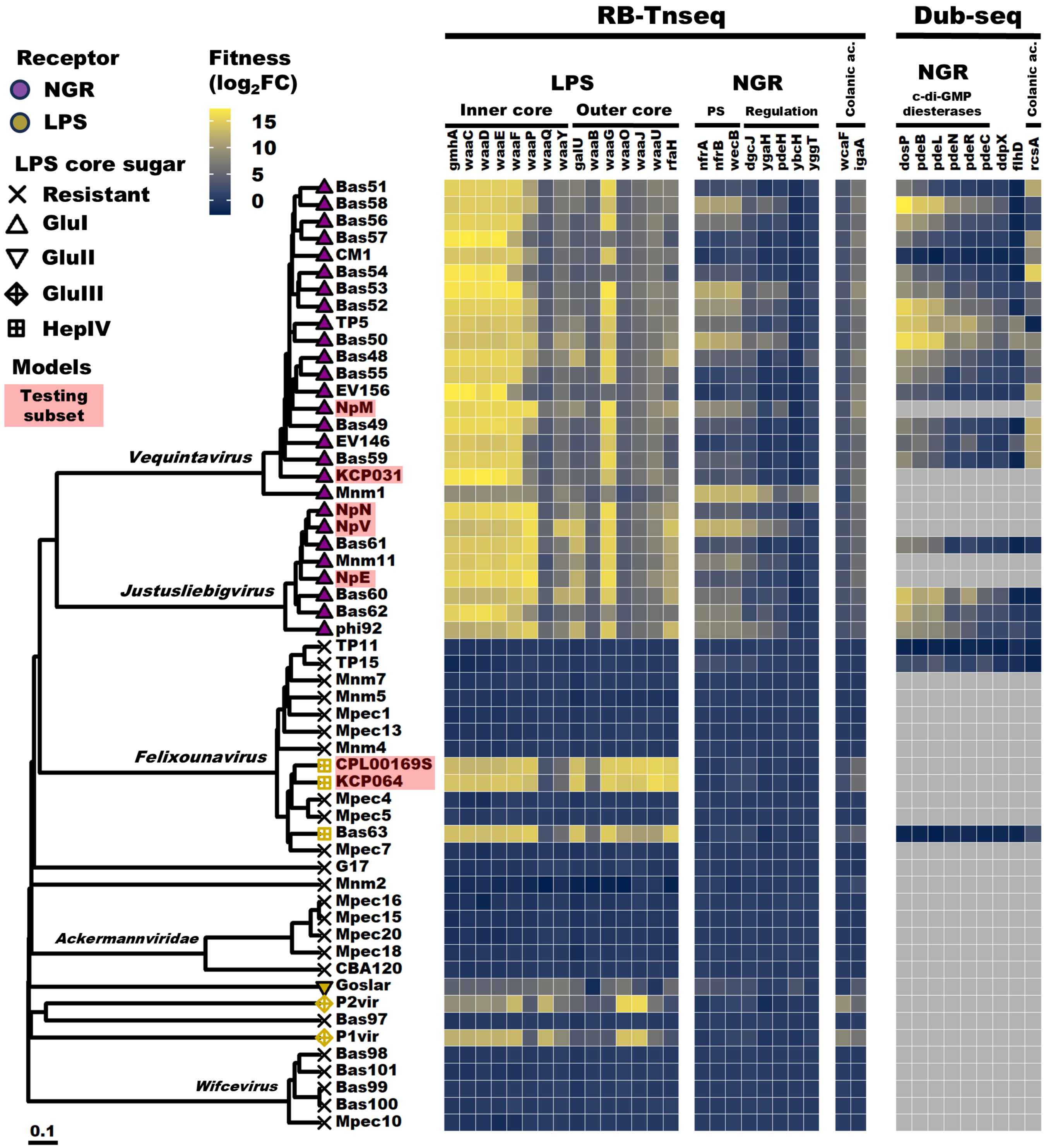


**Figure S4. RB-TnSeq-based receptor identification in other myoviruses.** Annotated heatmap showing the fitness values ($\log_2$ fold-change) for 34 high-scoring genes, the related membrane structures, and the resulting receptor(s) assignment in 56 non-*Straboviridae* myoviruses. Phages included in the model testing subset are highlighted in red. Grey boxes indicate missing data.

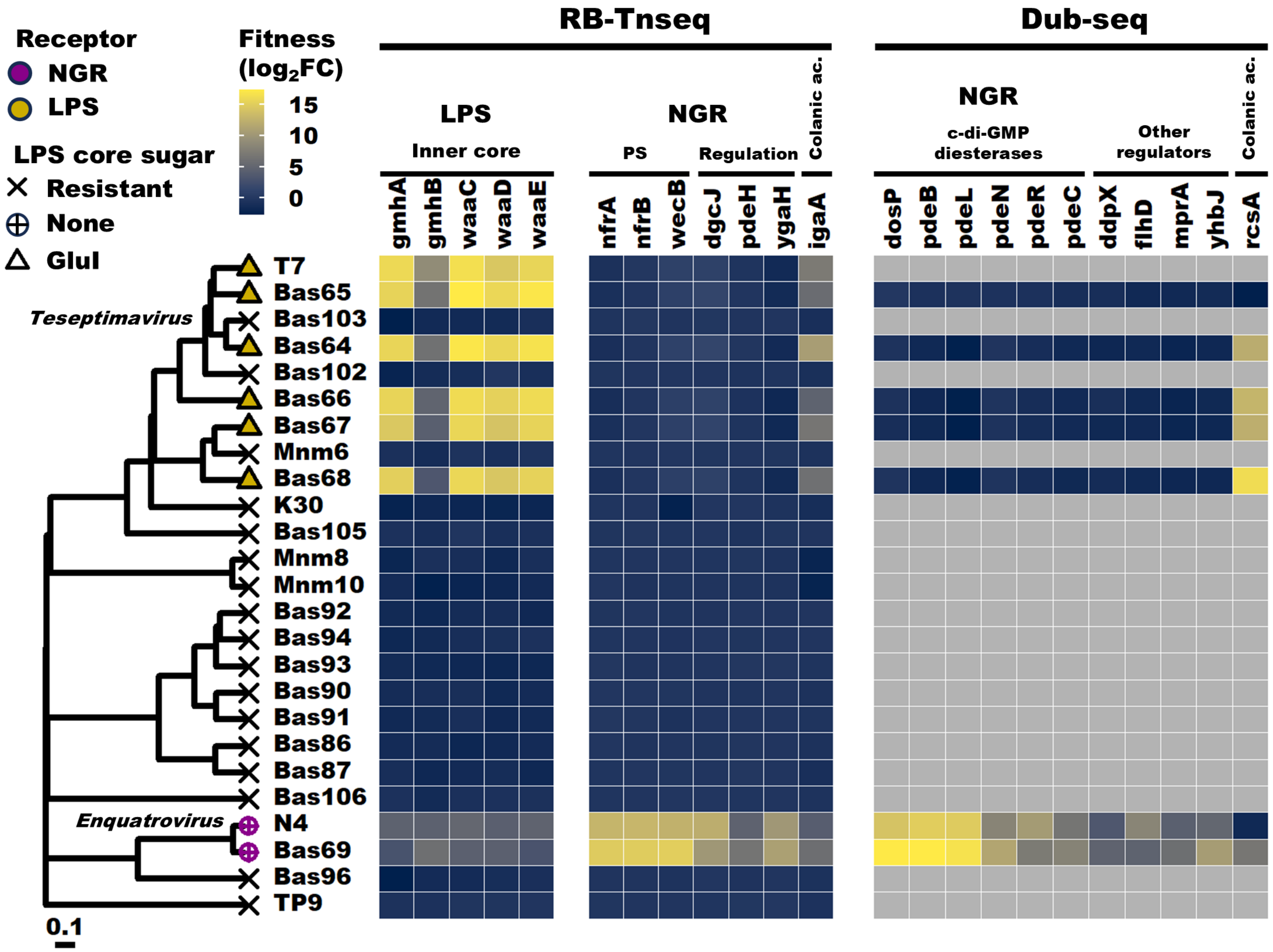


**Figure S5. RB-TnSeq-based receptor identification in podoviruses.** Annotated heatmap showing the fitness values ($\log_2$ fold-change) for 23 high-scoring genes, the related membrane structures, and the resulting receptor(s) assignment in 25 podoviruses. Grey boxes indicate missing data.

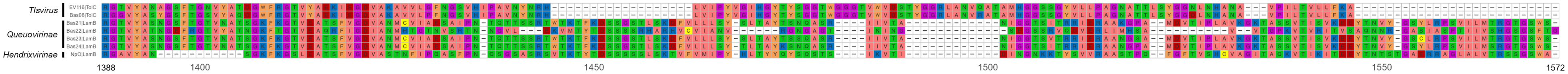


**Figure S6. TolC- and LamB-specific GpJ C-terminal domains identified by comparative genomics.** Protein alignment of the C-terminal domains of GpJ for 2 TolC- and 5 LamB-specific *Drexlerviridae* and related phages (alignment positions 1388 to 1572 / 1572 total). The central tail fiber GpJ of Lambda phage was not included as its C-terminal domain shared little to no homology with the 7 phages displayed. Relevant taxonomic levels and targeted receptors are indicated along the phage names. Amino acid residues are colored according to the “Zappo” color scheme.

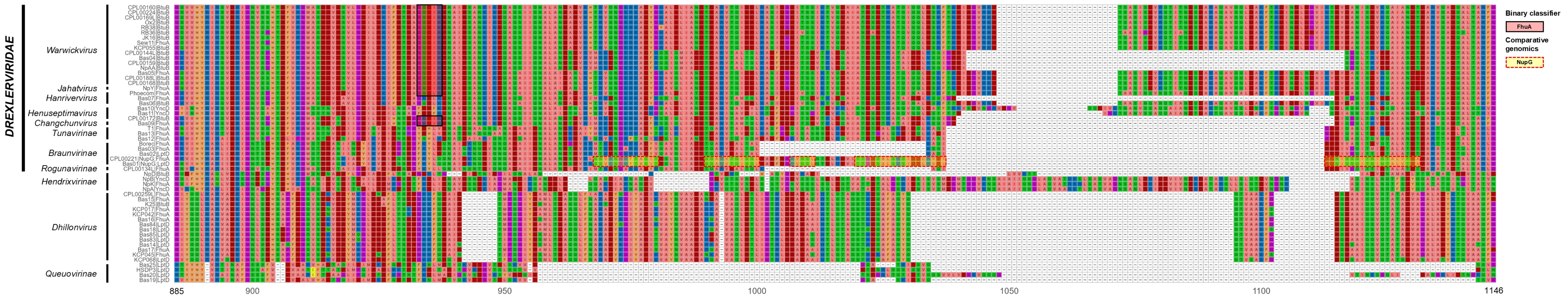


**Figure S7. NupG-specific GpJ C-terminal domains identified by comparative genomics.** Protein alignment of the central tail fiber GpJ C-terminal domains of 21 FhuA-, 18 BtuB-, 12 LptD-, 4 YncD-, and 2 NupG-specific *Drexlerviridae* and related phages (alignment positions 885 to 1146 / 1329 total). Relevant taxonomic levels and targeted receptors are indicated along the phage names. Predictive *k*-mer features identified by the FhuA binary classifier are highlighted by red boxes, and amino acid sequences identified by comparative genomics to be unique to the two NupG-specific phages (Bas01 and CPL00221) are highlighted by yellow boxes with dashed red outlines. Amino acid residues are colored according to the “Zappo” color scheme.

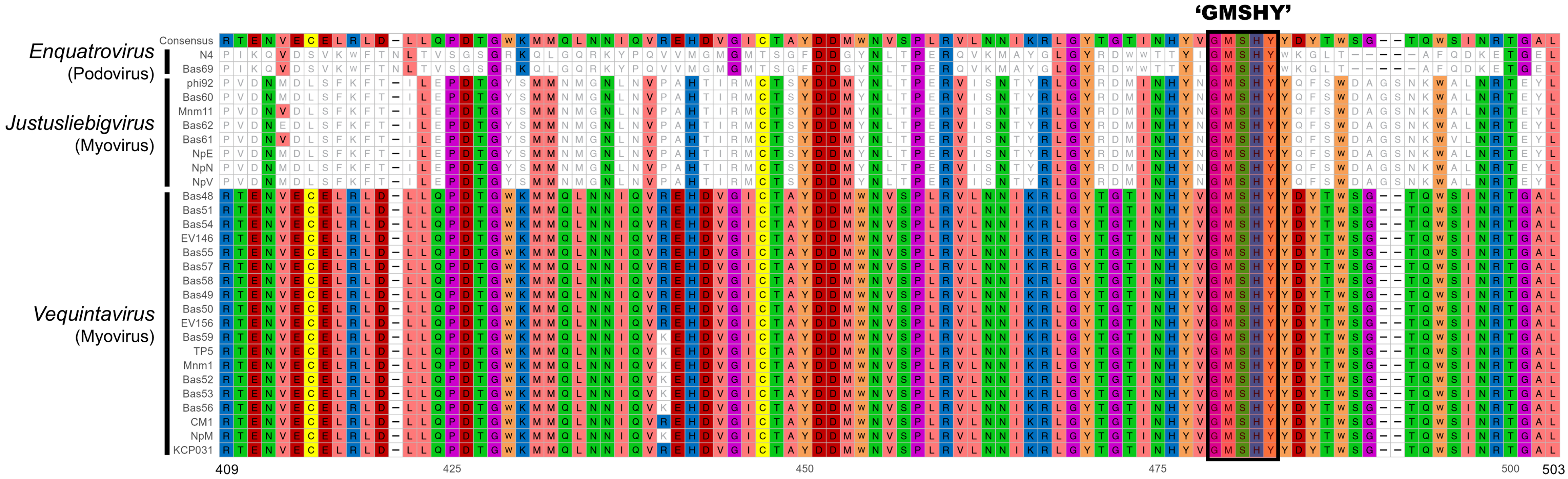


**Figure S8. NGR-specific homologous tail fiber domain identified by comparative genomics and annotation-free modelling.** Protein alignment of the putative NGR-binding homologous domain identified in the tail fiber genes of 27 myoviruses and 2 podoviruses (alignment positions 409 to 503 / 1504 total). Alignment is viewed in the “consensus = true” mode, i.e. every amino acid residue varying from the majority consensus is displayed in a white box with grey letters. Relevant taxonomic levels and morphotype information are indicated along the phage names. The predictive and perfectly conserved *k*-mer feature ‘GMSHY’ identified by the NGR binary classifier is highlighted with a black outline. Amino acid residues are colored according to the “Zappo” color scheme.

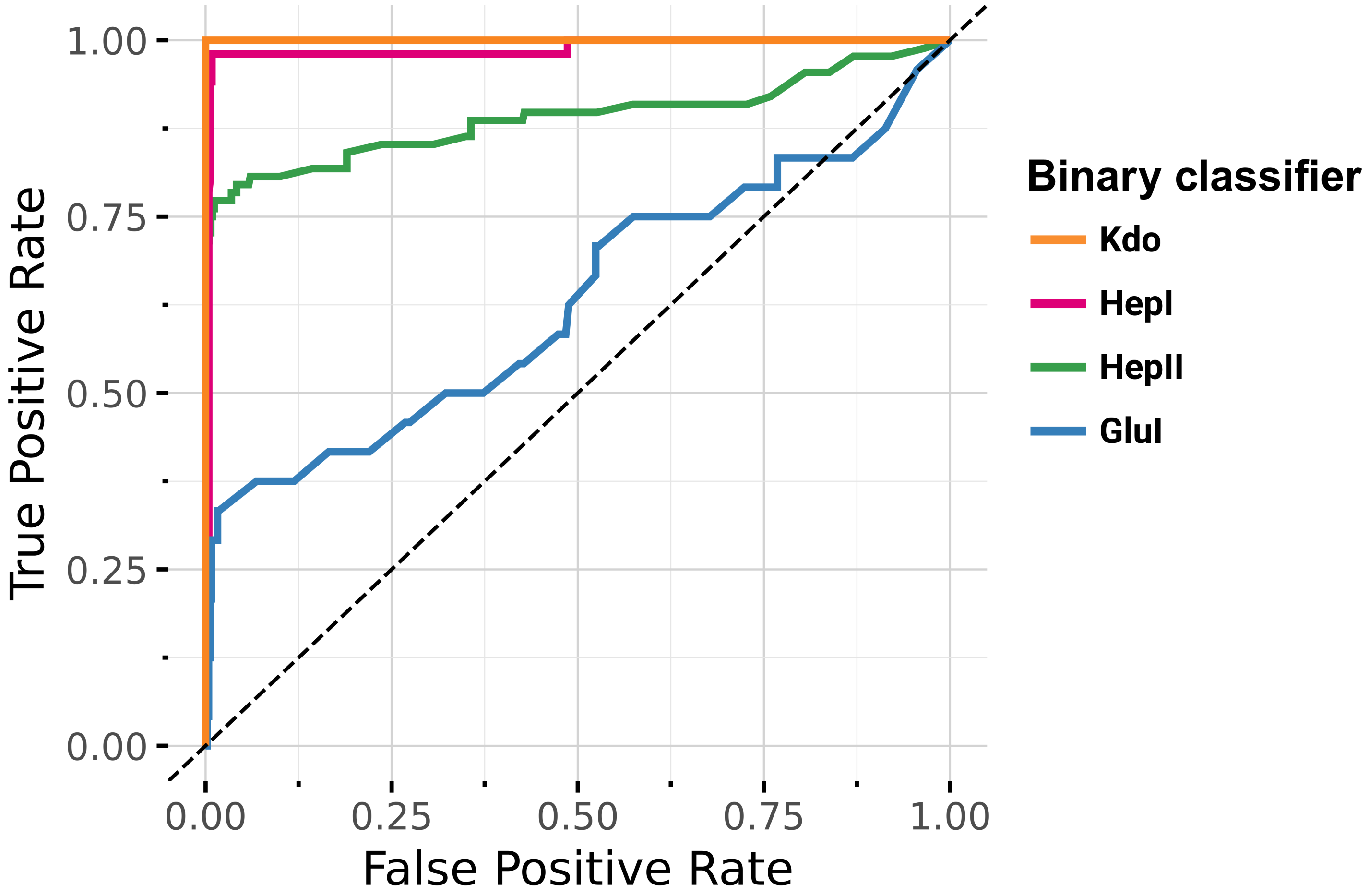


**Figure S9. ROC curves for the LPS core sugar binary classifiers.** Receiver operating characteristic (ROC) curves showing the sensitivity (true positive rate) and specificity (true negative rate) of the 4 LPS core sugar-specific binary classifiers (Kdo, HepI, HepII, GluI).

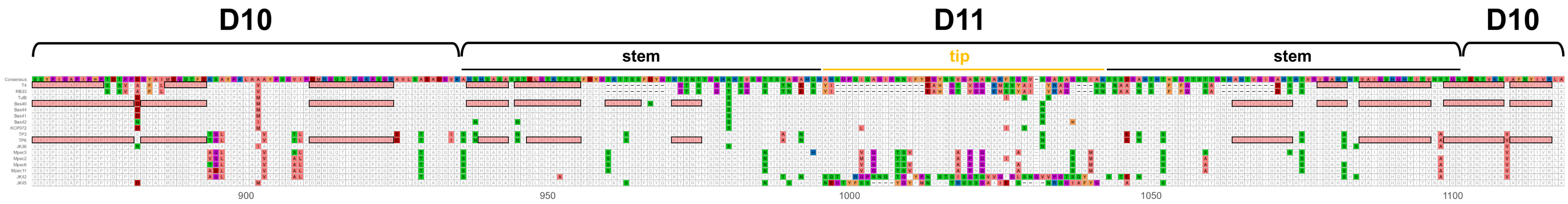


**Figure S10. OmpC-specific Gp37 C-terminal domains identified by annotation-free modelling.** Protein alignment of the Gp37 C-terminal domains of the 17 OmpC-specific *Straboviridae* phages (alignment positions 865 to 1118 / 1118 total). Alignment is viewed in the "consensus = false" mode, i.e. every amino acid residue varying from the majority consensus is displayed in colors. The structural domains D10 and D11, and the D11 "stem" and "tip" sub-domains described by Bartual *et al.* 2007 and Islam *et al.* 2019 are indicated above the alignment track. Contiguous predictive *k*-mers identified by the OmpC binary classifier are highlighted by red boxes for phages T4, Bas40, and TP6. Amino acid residues are colored according to the "Zappo" color scheme.

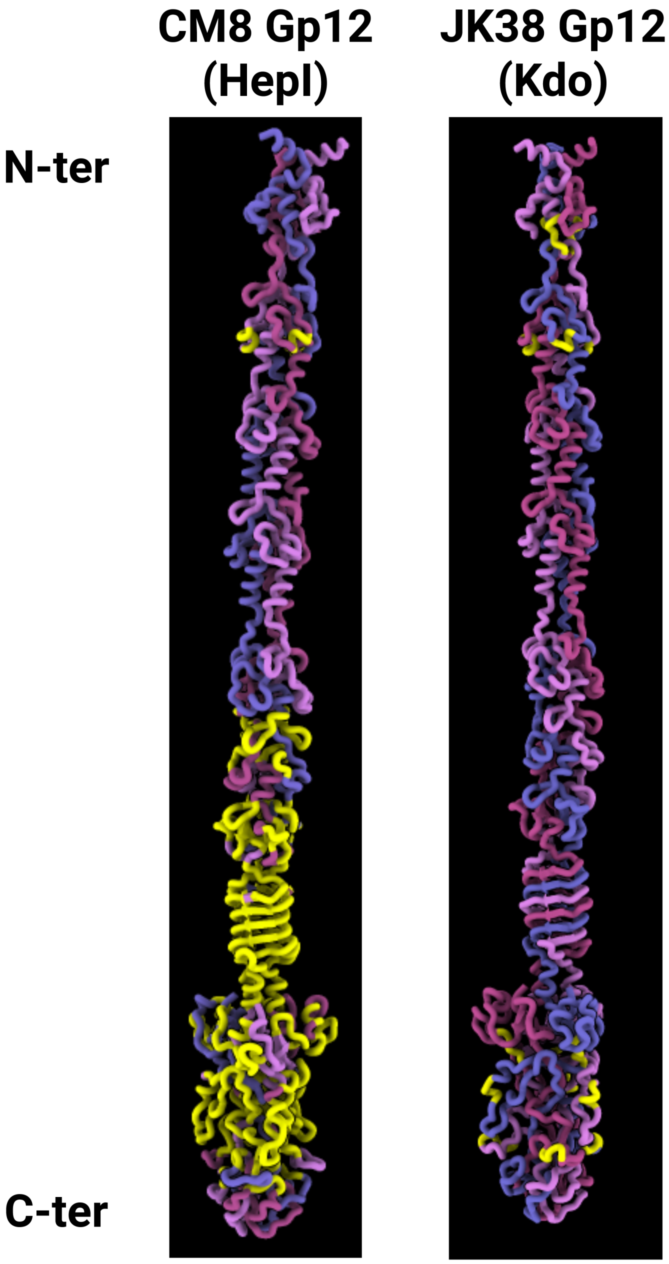


**Figure S11. AlphaFold3 models of representative instances of the Kdo- (JK38) and HepI-binding (CM8) Gp12 short tail fibers of *Straboviridae* phages.** Each model consists of a trimer of Gp12, and monomer chains are indicated by different colors. Contiguous predictive *k*-mers identified by the Kdo and HepI binary classifiers are highlighted on the models in yellow. The C-terminal domains at the bottom of the models are putatively interacting with the targeted LPS core sugars, while the N-terminal domains at the top are connected to the phage baseplate.

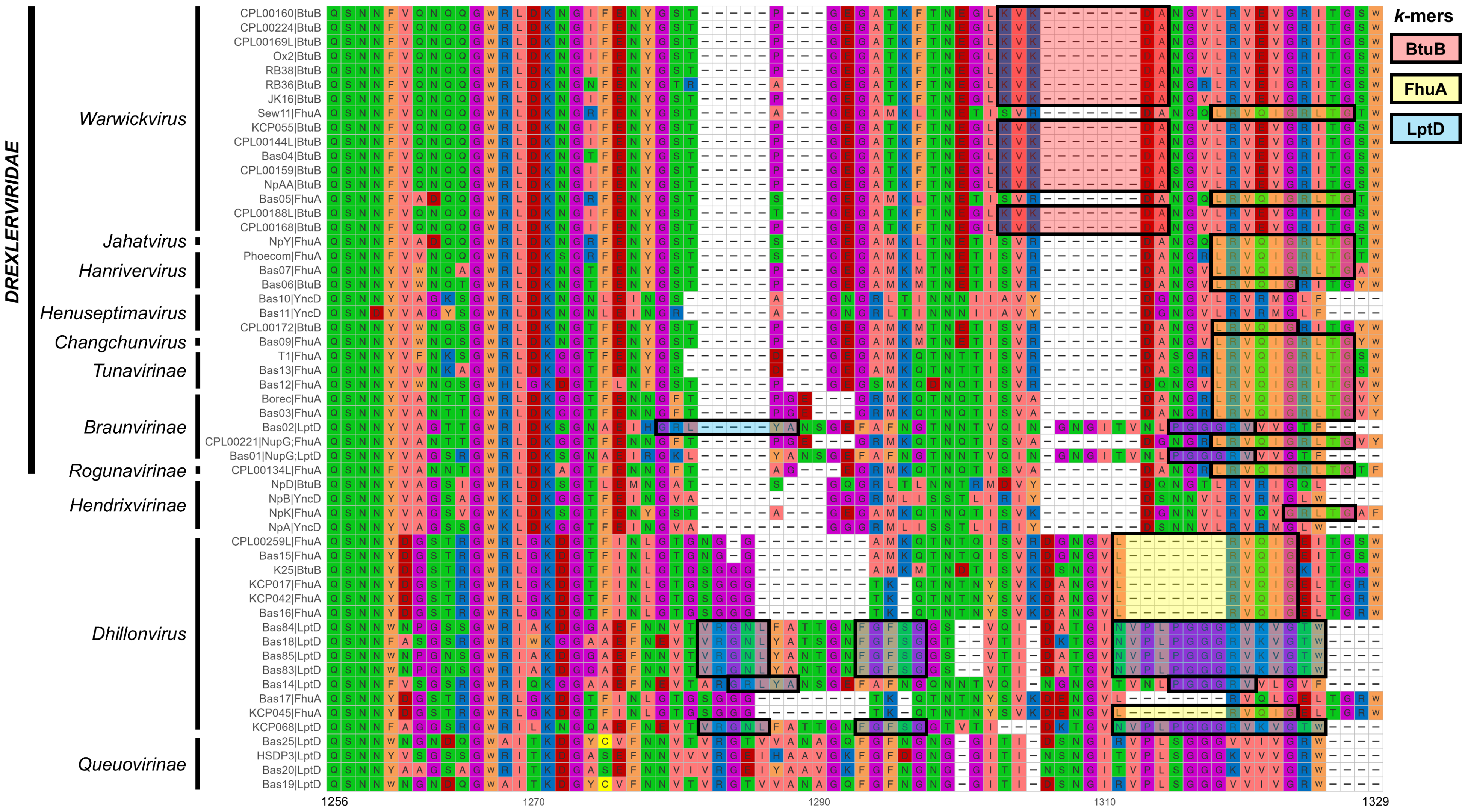


**Figure S12. BtuB-, FhuA- and LptD-specific GpJ C-terminal domains identified by annotation-free modelling.** Protein alignment of the central tail fiber GpJ C-terminal domains of 21 FhuA-, 18 BtuB-, 12 LptD-, 4 YncD-, and 2 NupG-specific *Drexlerviridae* and related phages (alignment positions 1256 to 1329 / 1329 total). Relevant taxonomic levels and targeted receptors are indicated along the phage names. Contiguous predictive *k*-mer features identified by the BtuB, FhuA, and LptD binary classifiers are highlighted by red, yellow, and blue boxes, respectively. Amino acid residues are colored according to the "Zappo" color scheme.

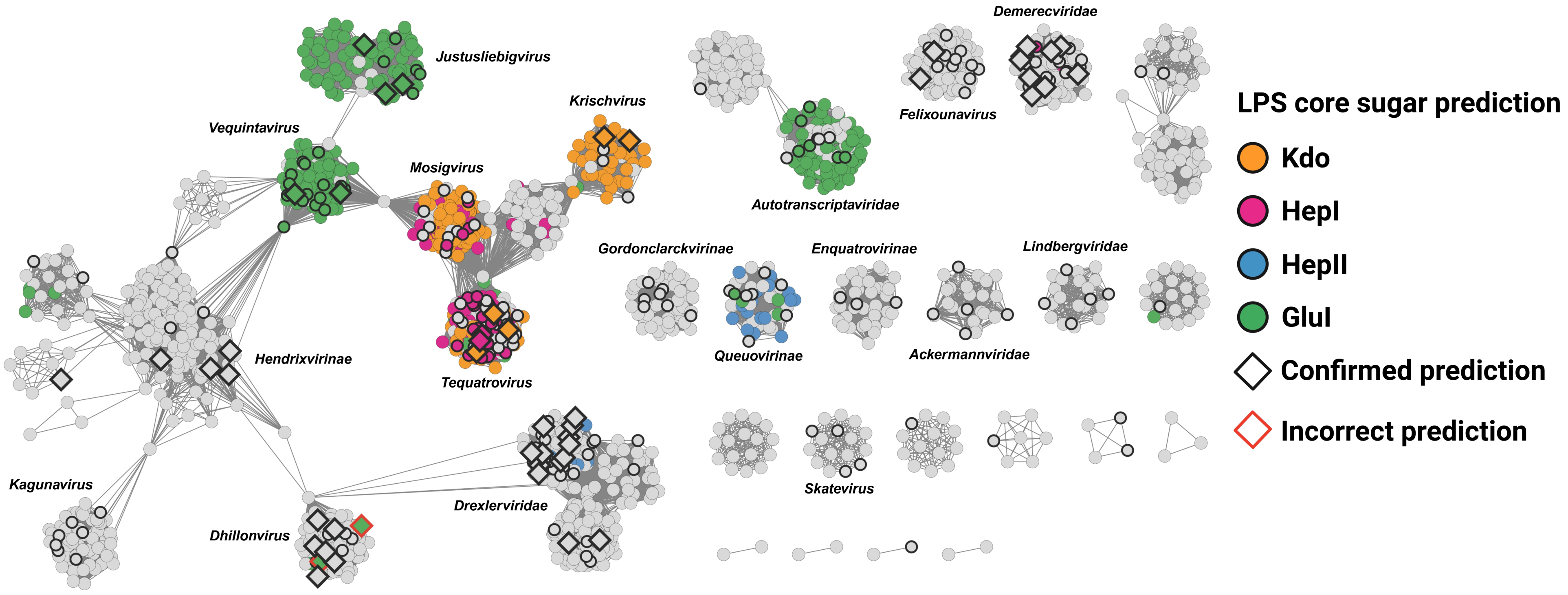


**Figure S13. LPS core sugar specificity predictions across NCBI *E. coli* phages.** vConTACT2 gene-sharing network displaying predictions of the 4 LPS core sugars binary classifiers (Kdo, HepI, HepII, and GluI) across 1,875 *E. coli* dsDNA phages from the NCBI GenBank database and the 255 phages in our collection. Phages included in the initial training set (206 phages) are represented with a circle, while phages selected for the testing subset (49 phages) are indicated by a diamond. Colors indicate the predicted receptor for NCBI phages or experimentally determined receptor for phages in the training and testing subsets.
